# Noncoding function of super enhancer derived mRNA in modulating neighboring gene expression and TAD interaction

**DOI:** 10.1101/2023.12.05.570115

**Authors:** Bingning Xie, Ann Dean

## Abstract

Super enhancers are important regulators of gene expression that often overlap with protein-coding genes. However, it is unclear whether the overlapping protein-coding genes and the mRNA derived from them contribute to enhancer activity. Using an erythroid-specific super enhancer that overlaps the *Cpox* gene as a model, we found that *Cpox* mRNA has a non-coding function in regulating neighboring protein-coding genes, eRNA expression and TAD interactions. Depletion of *Cpox* mRNA leads to accumulation of H3K27me3 and release of p300 from the *Cpox* locus, activating an intra-TAD enhancer and gene expression. Additionally, we identified a head-to-tail interaction between the TAD boundary genes *Cpox* and *Dcbld2* that is facilitated by a novel type of repressive loop anchored by p300 and PRC2/H3K27me3. Our results uncover a regulatory role for mRNA transcribed within a super enhancer context and provide insight into head-to-tail inter-gene interaction in the regulation of gene expression and oncogene activation.

## INTRODUCTION

Super enhancers are clusters of transcriptional enhancers that are collectively bound by groups of transcription factors and mediator, and play an important role in regulating cell identity genes(1). Some super enhancers overlap with protein-coding genes (1,2), but it is unclear whether these genes can function as enhancers themselves.

Controlling gene expression is a complex process that involves not only the direct interaction between an enhancer and its target gene, but also indirect influences from the expression status of other genes. For instance, the enhancer release and re-targeting (ERR) model proposes that repressing gene expression through transcription start site (TSS) deletion or dCas9-KRAB-based transcriptional repression can release its partner enhancer, which can then retarget to other neighboring genes, activating their expression(3). The involvement of mRNA in this process and the mechanism by which enhancer activation occurs in response to mRNA loss are currently unknown.

Loss of transcription by TSS deletion results in histone deacetylation, which mediates Ras-induced gene silencing and leads to accumulation of H3K27me3 in the gene body(4). PRC2 can interact with both RNA and chromatin, and they compete with each other for PRC2 binding(5). Nascent RNA- PRC2 interaction can inhibit PRC2 function in situ, as demonstrated by the long non-coding RNA LEVER, which is transcribed from upstream of the β-globin cluster and interacts with PRC2 in its nascent form. The application of CRISPR-Cas9-mediated knock-out or CRISPRi-based transcriptional inhibition of LEVER triggers PRC2 translocation to chromatin, leading to altered chromatin interaction between LEVER and its target sites and, subsequently, activation of neighboring ε-globin gene expression(6). However, it remains unclear whether the RNA molecule itself is involved in this process. Thus, currently, there is no direct evidence to support the hypothesis that H3K27me3 accumulation following TSS deletion is caused by PRC2 moving from RNA to chromatin.

The genome in the nucleus is organized into different layers of higher architecture, ranging from nucleosome to chromatin looping, to topologically associated domains (TADs), and A/B compartments(7,8). Some TAD boundaries overlap with protein-coding genes or non-coding genes (9-12). The non-coding RNA transcribed from a TAD boundary (bRNA) has been reported to strengthen TAD insulation by recruiting CTCF to the TAD boundary(9). R loops, which are DNA:RNA hybrids, form as natural byproducts of transcription and accumulate around transcription start and termination sites(13). They have been found to facilitate and stabilize chromatin loops and TADs (14-17). However, whether mRNA transcribed from TAD boundaries also has a regulatory role in TAD structure is currently unclear. Actively transcribed genes located at TAD boundaries have been reported to function as boundary elements. For example, the transcribed α2-globin gene, but not a nearby CTCF site, was the anchor for the TAD boundary in the α-globin locus(11). Nevertheless, it is not known whether genes located at the two boundaries of a TAD regulate each other.

In this study, we demonstrate that *Cpox*, which overlaps with a super enhancer located at a TAD boundary, transcribes a mRNA that has a non-coding function in regulating enhancer and neighboring gene expression as well as intra-TAD interactions. Intra-TAD protein coding gene *St3gal6* and enhancer *CpoxeRNA* interact with the gene body region of *Cpox*. The two TAD anchor genes, *Cpox* and *Dcbld2*, are organized in a head-to-tail inter-gene conformation. A novel type of repressive loop established by a p300 only peak in the *Cpox* intron and H3K27me3 in the promoter of *Dcbld2*, represses *Dcbld2* gene expression. Upon loss of *Cpox* mRNA, PRC2 translocation from mRNA to chromatin alters these chromatin interactions and releases transcription activators from the *Cpox* locus, resulting in the activation of the enhancer and protein coding genes *St3gal6* and *Dcbld2* within the TAD. This, in turn, leads to increased transcriptional R-loops and strengthens TAD corner loop interaction. Analysis of cancer mutations shows that the head-to-tail TAD boundary gene conformation is a potential mechanism to harness oncogene activation.

## MATERIALS AND METHODS

### Cell culture

MEL cells were cultured in Dulbecco’s modified Eagle’s medium supplemented with 10% fetal bovine serum in a humidified incubator with 5% CO2. Differentiation of MEL cells was induced by 2% Dimethylsulfoxide for 5 days.

### Knock down experiments: shRNA and ASO

shRNA knock down experiments were performed by electroporation of shRNA (sigma MISSION® shRNA) into MEL cells with BTXpress High Performance Electroporation Kit & Solution (BTXpress, catalog no. 89130-538) and Biorad instrument. Cells were transferred to DMEM medium containing 20% FBS immediately after electroporation and incubated at 37 °C for two days, then cultured in DMEM medium with 10% FBS and 2μg/ml puromycin (Thermofisher, catalog no. A1113803) for 3 days. For shRNA knock down in 293T cell, two shRNAs for each gene (except one shRNA for *PSMD3*) were transiently transfected to 293T cell with lipofectamine 2000 (ThermoFiser, 11668019), selected with 2 μg/ml puromycin then collected for RNA isolation.

ASO knock down experiment was performed by electroporation of LNA(QIAGEN) into MEL cells with BTXpress High Performance Electroporation Kit & Solution (BTXpress, catalog no. 89130-538). Cells were transferred to DMEM medium containing 20% FBS immediately after electroporation and incubated at 37 °C for two days, then harvested for RNA isolation or incubated for one day before addition of 2% DMSO for differentiation for 4 days.

### CRISPR-Cas9 knock out

sgRNAs were designed by using CRISPick (Broad institute, MIT), cloned into pSpCas9(BB)-2A-Puro (PX459) V2.0 (pSpCas9(BB)-2A-Puro (PX459) V2.0 was a gift from Feng Zhang (Addgene plasmid # 62988; http://n2t.net/addgene:62988; RRID: Addgene_62988)) according to the protocol from Ran F., et. al(18). For *Cpox* intron 5 TFBS deletion, sgRNA plasmids were electroporated into MEL cells with BTXpress High Performance Electroporation Kit & Solution (BTXpress, catalog no. 89130-538). Cells were transferred immediately into DMEM medium with 20% FBS and cultured for one day, then treated with 4μg/ml puromycin for 12 hours, washed and cultured in puromycin-free DMEM with 10% FBS. *Cpox* TSS deletion was performed with same procedure except continuous cultured in medium with 2μg/ml puromycin to select permanent integrated cells. Selection of single colonies by serial dilution culture was performed in 96 well plates. Genotyping of single colonies was performed after extraction of DNA with QuickExtract DNA Extraction Solution (Biosearch Technologies, catalog no. 76081-766) and PCR amplification with Q5® High-Fidelity 2X Master Mix (NEB, catalog no. M0492S). PCR products were confirmed by Sanger sequencing (Genewiz)

### dCasRX

sgRNAs were designed by using cas13design, cloned into pXR003: CasRx gRNA cloning backbone (pXR003: CasRx gRNA cloning backbone was a gift from Patrick Hsu (Addgene plasmid # 109053; http://n2t.net/addgene:109053; RRID:Addgene_109053)), then co-electroporated with pXR002: EF1a-dCasRx-2A-EGFP (pXR002: EF1a-dCasRx-2A-EGFP was a gift from Patrick Hsu (Addgene plasmid # 109050; http://n2t.net/addgene:109050; RRID:Addgene_109050))(19) into MEL cell. Transfected cells were collected for extracting RNA 48h after electroporation.

### Flow Cytometry Assays

Flow cytometry experiments were conducted on FACS BD LSRFortessa machine (BD Biosciences). Cells were immunostained with PE Rat Anti-Mouse CD71 (BD Biosciences, Catalog #:553267), allophycocyanin-conjugated anti-TER119 (BD Biosciences, Catalog #:557909) antibodies, and 5 μg/mL Hoechst 33342 (Thermofisher, Catalog #: 62249)(20,21).

### Nascent RNA

Nascent RNA was detected by EU labeling experiments. Experiment was done according to the protocol of Click-iT Nascent RNA Capture kit (Invitrogen). Briefly, cells were pulsed with 0.5 mM EU for 1 hour, total RNA was isolated and 5 μg EU labeled RNA were used in a copper catalyzed click reaction with azide-modified biotin. Then 1μg biotinylated RNA were captured on streptavidin magnetic beads and cDNA synthesis performed with pull downed material directly on the beads using Superscript VILO cDNA synthesis kit (Invitrogen) followed by analysis with qRT-PCR. See Supplementary Data for primers.

### RT-qPCR

RNA was isolated with RNeasy Plus Mini Kit (QIAGEN, catalog no. 74134). 2 μg RNA was reverse transcribed into cDNA by SuperScript™ III First-Strand Synthesis System (Thermofisher, catalog no.18080051), and then analyzed by qPCR with iTaq Universal SYBR Green Supermix (Biorad, catalog no. 1725124) using a QuantStudio 6 FLEX machine (Thermofisher). See Supplementary Data for primers. Actin was used as internal control gene. Data were analyzed with -ΔΔCt method for Fig. 1C, -ΔCt method for Figure 3F and supplementary Fig. 1F, and 2^-ΔΔCt^ method for the rest of the data(22,23).

**Figure 1.**
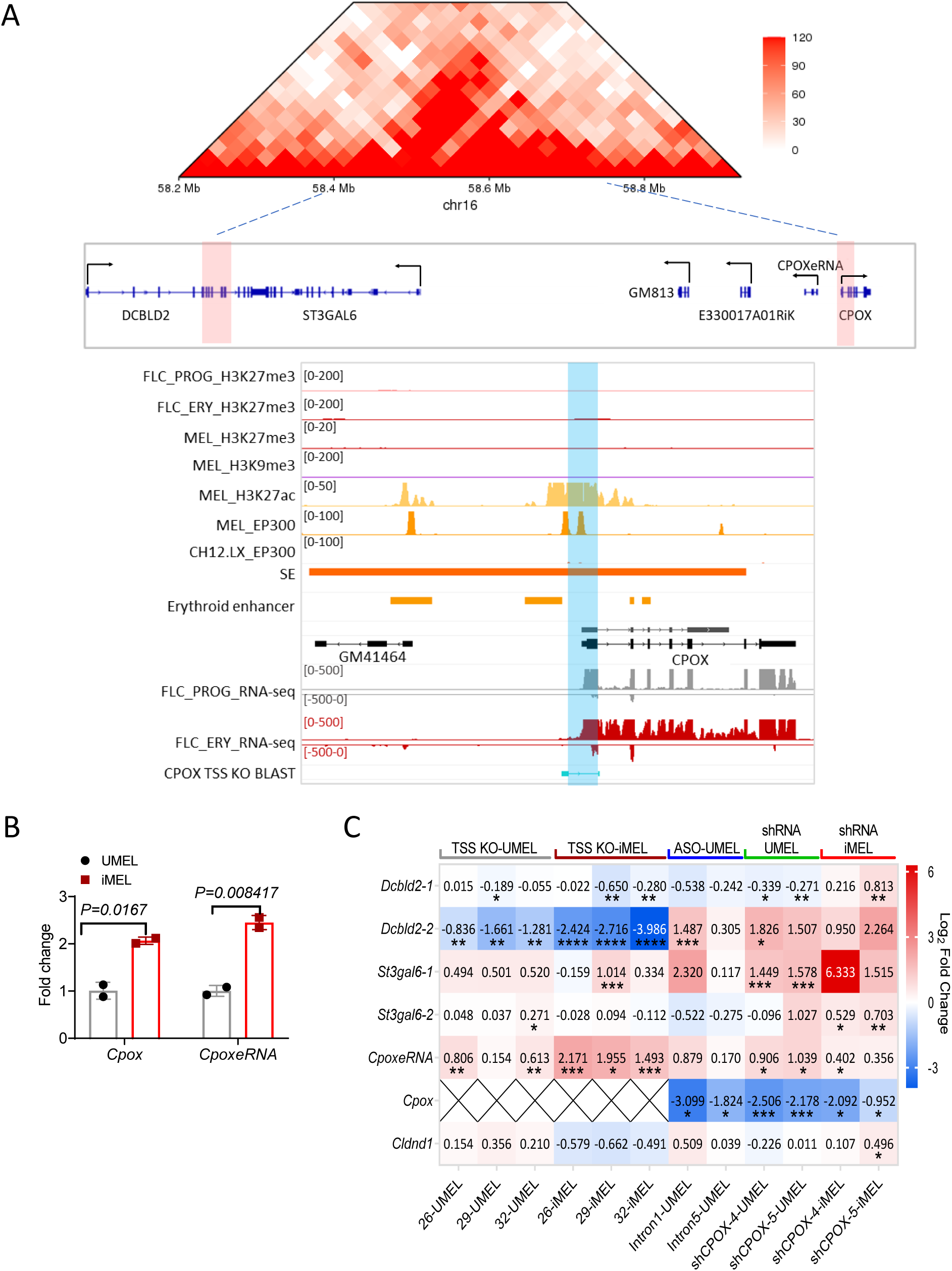
mRNA loss activates partner enhancer and neighboring genes. A. 10kb resolution HiC map of *Cpox* TAD from E14.5 Fetal liver cells (upper panel). Genes located in this TAD are shown below, TAD boundaries are highlighted in red, black arrows represent transcription direction. IGV tracks show the super enhancer overlap with *Cpox*, erythroid enhancers, histone ChIP-seq and RNA-seq data from progenitor (FLC_PROG) and differentiated mouse fetal liver cells (FLC_ERY), MEL cells and CH12.LX cells. *Cpox* TSS deleted region is highlighted in blue. **B.** qRT-PCR result shows *Cpox* and *CpoxeRNA* expression level in undifferentiated (UMEL) and differentiated (iMEL) MEL cells. Two biological replicates. Data are mean ± s.d., unpaired two-tailed t-test. **C.** Heatmap shows the qRT-PCR result of eRNA and gene expression levels in *Cpox* TSS deletion undifferentiated and differentiated MEL cells, *Cpox* knock down with ASO in undifferentiated MEL cells, and *Cpox* knock down with shRNA in undifferentiated and differentiated MEL cells. Value in each cell represent log2 fold change relative to corresponding control samples, data normalize with ACT. *: P ≤ 0.05; **: P ≤ 0.01, ***: P ≤ 0.001, ****: P ≤ 0.0001, unpaired one-tailed t-test.

### ChIP-qPCR

MEL cells differentiated 5 days in 2% DMSO were used for Chromatin immunoprecipitation as described(24). Eluted DNA was analyzed by qPCR with iTaq Universal SYBR Green Supermix (Biorad, catalog no. 1725124) and QuantStudio 6 FLEX machine (Thermofisher). Antibodies used for immunoprecipitation were: EZH2 (Cell Signaling, 5246), H3K27me3 (Cell Signaling, 9733), H3 (Abcam, ab1791). For p300 ChIP experiment, undifferentiated MEL cells were used, p300 antibody (Santa Cruz Biotechnology, sc-48343). See Supplementary Data for primers.

### fCLiP-qPCR

MEL cells differentiated for 5 days in 2% DMSO were used for fCLiP as described(25). Eluted RNA was reverse transcribed into cDNA with SuperScript™ III First-Strand Synthesis System (Thermofisher, catalog no.18080051), then analyzed by qPCR with iTaq Universal SYBR Green Supermix (Biorad, catalog no. 1725124) and QuantStudio 6 FLEX machine (Thermofisher). Antibodies used for immunoprecipitation were: EZH2(Cell Signaling, 5246), SUZ12(Cell Signaling, 3737). See Supplementary Data for primers.

### 3C-qPCR

MEL cells differentiated for 5 days in 2% DMSO were used for 3C experiments performed as described(26,27). DNA was analyzed by qPCR with iTaq Universal SYBR Green Supermix (Biorad, catalog no. 1725124) and QuantStudio 6 FLEX machine (Thermofisher). BAC controls were obtained from BACPAC company; alpha Aortic Actin-2(RP23-2N15), *Dcbld2*-*Cpox* locus (RP23- 317L24, RP24-264K24, RP24-371018). See Supplementary Data for primers.

### DRIP-qPCR

Undifferentiated MEL cells were used for S9.6(KeraFAST, ENH001) immunoprecipitation performed as described(14,28). Eluted DNA was analyzed by qPCR with iTaq Universal SYBR Green Supermix (Biorad, catalog no. 1725124) and QuantStudio 6 FLEX machine (Thermofisher). See Supplementary Data for primers.

### Western blot

Protein was extracted with RIPA buffer in the presence of Halt™ Protease Inhibitor Cocktail (Thermofisher, catalog no.78443) and quantified with Pierce™ BCA Protein Assay Kit (Thermofisher, catalog no. 23227). 20 μg protein was use for western blotting. Antibodies used were: EZH2(Cell Signaling, 5246), H3K27me3 (Cell Signaling, 9733), ACT (Sigma, A3853), CPOX (Santa Cruz Biotechnology, sc-393388), a-Tubulin (Santa Cruz Biotechnology, sc-5286).

### GSK343 treatment

Undifferentiated shCTRL and shCPOX-4 Cells seeded at 0.5M cells/ml and treated with 5 μl 10mM GSK343 (Sigma, catalog no. SML076, final concentration 5μM) or 5μl DMSO (vehicle) for 24 hours.

### Data Analysis

#### ChIP-seq data analysis

ChIP-seq data are all from ENCODE. Bedtools(29) slop was used to generate the extended H32K7ac region (+/-1kb around H3K27ac peak) file. Pybedtools(30) was used to find the overlapping region between p300 and extended H3K27ac region(+/-1kb around H3K27ac peak). Heatmap for H3K27ac, H3K27me3, and p300 and binding profile of MYC, GATA2, and ELF1 around p300 only peaks were analyzed by deeptools (v. 3.5.1)(31). Pie chart of p300 only peak genome distribution was analyzed by ChIPseeker (v. 1.18.0)(32). Motif analysis were performed with HOMER(33).

#### Hi-C data analysis and generation of virtual 4C profiles

Contact matrix files(.hic) were obtained from the 4DN data portal(34) for Late G1 phase G1ER HiC(4DNFI6H926RO) and CH12.LX(4DNFI8KBXYNL). Figure 1A Fetal liver HiC raw Fastq files were obtained from(35), raw sequence data was mapped and processed using Juicer(36). HiC matrix and virtual 4C were analyzed by using GENOVA(dev version)(37). Supplementary Fig. 6A and D HiC matrices were displayed by Juicebox(38). Supplementary Fig. 2D, 6A and 6D insulation score and boundary track were obtained from 4DN data portal. G1E-ER4 and CH12X RNA seq data were obtained from ENCODE. Supplementary Fig. 1G processed HiC matrix was download from GSE184974(39), *Dcbld2*-*Cpox* TAD HiC map was analyzed with GENOVA(dev version)(37).

#### Interaction Categories, aggregate peak analysis and loop/TAD gene expression (Fig. 5)

Loop and TAD files are obtained from G1ER cell cycle in situ HiC (GSE129997)(40). For TAD, +/-5kb region of each boundary was used for further analysis. Files from each cell cycle were merged by using HiCExplorer hicMergeLoops(41). Interaction categories (Fig. 5E) were identified with GenomicInteractions(42), promoter was defined as +/- 1kb around transcription start site while terminator was defined as +/- 5kb around the end of an annotated transcript. Lollipop plot was draw with home-made R script. Fig. 5F and G aggregated peak analysis was done by using Juicer (v. 1.6)(36), with 25kb bins and window 6, normalized with VC_SQRT. G1ER in situ HiC late G1 phase .hic file (<u>4DNFI6H926RO</u>) was from 4DN database(34). Fig. 5H gene expression data was obtained from ENCODE (ENCFF199TJO.tsv) and was analyzed with home-made R script and plotted with ggpubr package.

#### TAD boundary interaction categories, gene expression, and oncoplot (Fig. 6 and supplementary Fig.S6 and S7)

TAD files are obtained from Rao S. et al., 2014 (GSE63525)(8), +/-5kb region was used as TAD boundary for further analysis. Interaction categories (Fig. 6A and Supplementary Fig. 6A) were identified with GenomicInteractions(42), and Bar plot was drawn with ggpubr. Fig. 6B and Supplementary Fig. 6B, gene expression data was obtained from ENCODE and was analyzed with home-made R script and plotted with ggpubr package. Supplementary Fig. 7B, mutation at TSS was analyzed with ICGC release 28 data by using script from Oh et al(3) and Oncoplot was drawn with maftools(43).

### Statistics and reproducibility

Statistical analysis was carried out using both GraphPad Prism 9 Software and t.test function in excel. Figure 5H, Figure 6B, and Supplementary Fig. 6B were using R and detailed statistical methods are annotated in the figure legend. The types of the statistical tests and the exact value of “n” are reported in the figure legends.

## RESULTS

### Depletion of super enhancer derived mRNA activates enhancer and neighboring genes

The *Cpox* gene encodes the sixth enzyme in the heme synthetic pathway. HiC experiment using murine fetal liver erythroid cells show that *Cpox* is situated at the right boundary of a TAD that contains protein-coding genes *Dcbld2* and *St3gal6*, as well as non-coding genes *Gm813*, *Gm49701*, and *E330017A01Rik*(35) (Fig. 1A). The *Dcbld2* gene, which has multiple isoforms, is located at the left TAD boundary. The promoter of the long isoform, *Dcbld2*-1, is situated outside the TAD boundary, while the promoter of the short isoform, *Dcbld2*-2, overlaps with the TAD boundary. According to annotation data from the dbSUPER super enhancer database(44), *Cpox* overlaps with a super enhancer that incorporates four erythroid specific enhancers(45) (Fig. 1A). Notably, the 5’ end of the super enhancer can be transcribed into an enhancer RNA. We named this RNA molecule *CpoxeRNA*. Both *Cpox* mRNA and *CpoxeRNA* are upregulated during erythropoiesis, and *CpoxeRNA* shows an erythroid specific expression pattern (Fig. 1B and Supplementary Fig. 1A).

To dissect the function of the protein-coding gene within the super enhancer, we deleted the transcription start site (TSS) of *Cpox* in MEL cells (supplementary Fig. 1B-D). After *Cpox* TSS deletion, *St3gal6* and *CpoxeRNA* were up-regulated and *Dcbld2-2* was down-regulated in both undifferentiated and differentiated cells, while *Dcbld2-1* was only decreased in differentiated cells (Fig.1C, and Supplementary Fig. 1E for primer location for *Dcbld2* and *St3gal6*). We next conducted experiments to knock down (KD) *Cpox* mRNA using two different methods. First, we employed shRNAs targeting exon 5 (shCPOX-4) or exon 7 (shCPOX-5) of *Cpox*. In undifferentiated MEL cell (UMEL), *CpoxeRNA*, *St3gal6-1* and *Dcbld2*-2 were up regulated by *Cpox* mRNA KD, while *Cldnd1*, which is located outside of this TAD, was not affected (Fig. 1C). In differentiated MEL cell (iMEL) with *Cpox* KD, *St3gal6* and *CpoxeRNA* were upregulated, as well as both isoforms of *Dcbld2*. This result was confirmed by a different knockdown method using anti-sense oligonucleotides (ASO) to target intron 1 or intron 5 of *Cpox* (Fig. 1C). We observed a shift toward the predominant expression of the long isoform of *Dcbld2* (*Dcbld2*-1) in differentiated wild type MEL cells (iMEL) (see supplementary Fig. 1F). This observation may account for the discrepancy in *Dcbld2*-1 expression between UMEL and iMEL cells following *Cpox* mRNA knockdown. The discrepancy of *Dcbld2* expression between *Cpox* TSS deleted cells and *Cpox* mRNA knockdown cells suggest that the CPOX genomic locus which overlapping with the super enhancer, functions as the enhancer for *Dcbld2* and indicates that the intact TAD structure is essential for *Dcbld2* expression. The TAD corner interaction slightly increased after differentiation, coinciding with the upregulation of the long isoform of *Dcbld2* (*Dcbld2-1*) (supplementary Fig. 1F and G). Notably, depletion of *Cpox* mRNA did not affect erythropoiesis as judged by production of hemoglobin and erythroid cell surface markers upon differentiation (Supplementary Fig. 2 A, B). This observation is consistent with a previous study(46) reporting that *Cpox W373X* mutant creates a premature stop codon and results in reduced mRNA level and protein level. Homozygous *Cpox^W373X/W373X^* mice exhibited embryonic lethality at E9.5 day although a vascularized yolk sac with evidence of blood cell formation was observed. These findings support that *Cpox* absence can be bypassed in erythropoiesis. Our qRT-PCR analysis of nascent RNA in undifferentiated MEL cell confirmed that the upregulation of *Dcbld2*, *St3gal6*, and *CpoxeRNA* following *Cpox* mRNA knockdown was due to increased transcription (Supplementary Fig. 2C). Our results suggest that the mRNA transcribed from *Cpox* within a super enhancer has a non-coding function in regulating the expression of neighboring genes. Furthermore, our findings provide evidence that the mRNA transcribed from the super enhancer can act as a repressor, while its genomic locus functions as an enhancer.

### *Cpox* mRNA knock down affects intra-TAD interactions

To explore the effect of *Cpox* mRNA knock down on intra-TAD interactions, we first focused on the only protein coding gene located inside of this TAD— *St3gal6*. We performed a 3C experiment using the *St3gal6* promoter as anchor, since 3C has the highest resolution and is without bias present in capture based techniques. We found the *St3gal6* promoter mainly interacts with the *Cpox* gene body 1 region, which overlaps with the core region of the TAD boundary. The interactions between the *St3gal6* promoter and *Cpox* promoter/gene body regions are all modestly reduced in the *Cpox* mRNA knock down cells (Fig. 2A).

**Figure 2.**
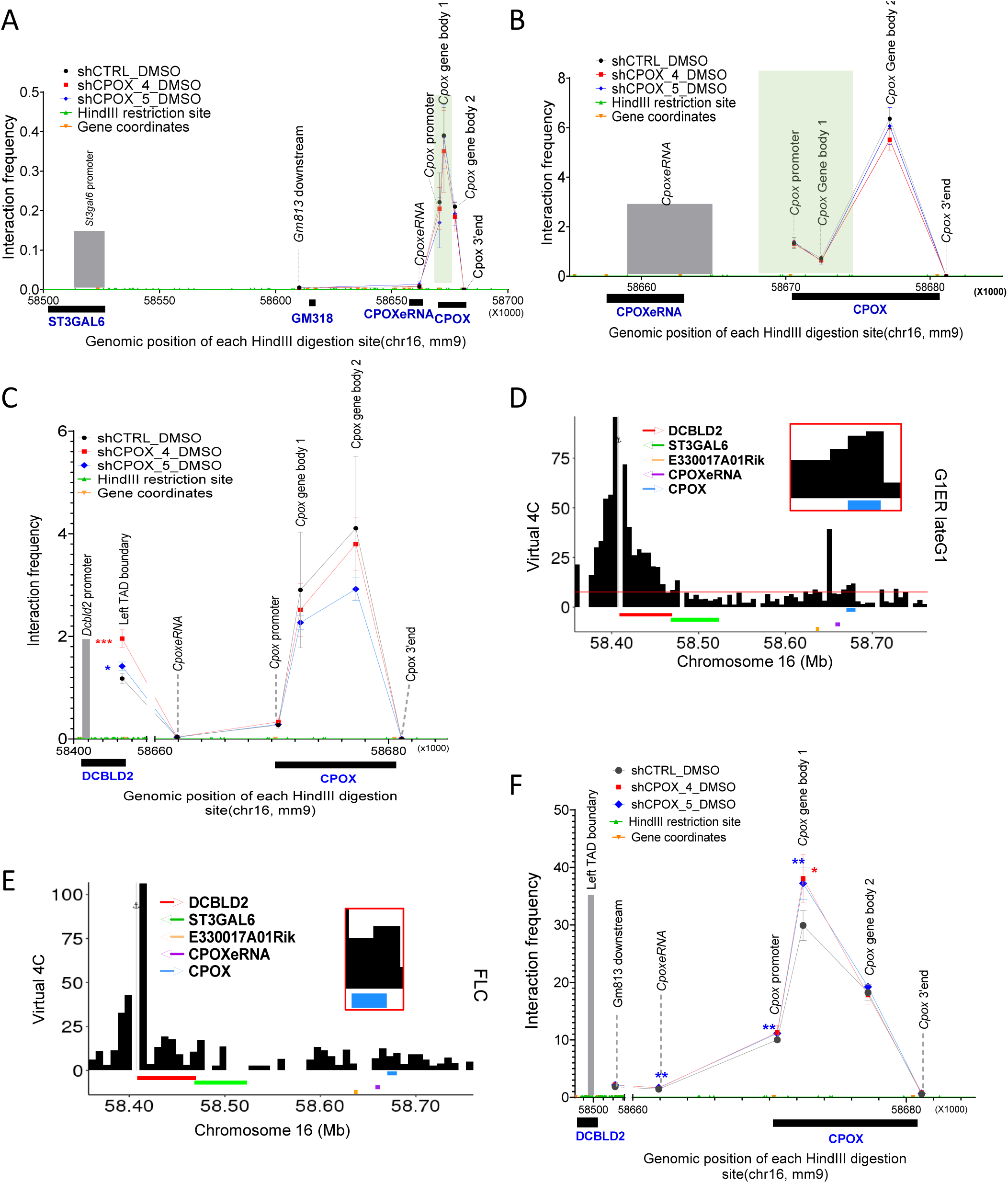
Chromatin interaction change after *Cpox* knock down. A. 3C-qPCR result shows the interaction of *St3gal6* promoter with *CpoxeRNA* and *Cpox* locus in control and *Cpox* knock down cells, *St3gal6* promoter as anchor. TAD boundary regions are highlighted in green. Three biological replicates. Data are mean ± s.d., *: P ≤ 0.05; **: P ≤ 0.01, ***: P ≤ 0.001, ****: P ≤ 0.0001, unpaired two-tailed t-test. **B.** 3C-qPCR result shows the interaction of *CpoxeRNA* with *Cpox* locus in control and *Cpox* knock down cells. *CpoxeRNA* locus as anchor. TAD boundary regions are highlighted in green. Three biological replicates. Data are mean ± s.d., *: P ≤ 0.05; **: P ≤ 0.01, ***: P ≤ 0.001, ****: P ≤ 0.0001, unpaired two-tailed t-test. **C.** 3C-qPCR result shows the interaction of the *Dcbld2* promoter with the *Cpox* 3’ region, *Dcbld2* promoter as anchor. TAD boundary regions are highlighted in green. Four biological replicates for shCTRL, three biological replicates for shCPOX- 4 and shCPOX-5. Data are mean ± s.d.. *: P ≤ 0.05; **: P ≤ 0.01, ***: P ≤ 0.001, ****: P ≤ 0.0001, unpaired two-tailed t-test. **D.** Virtual 4C plot from G1ER late G1 phase in situ HiC data shows the interaction between the *Dcbld2* promoter and *Cpox* 3’ region. *Dcbld2* promoter as anchor. **E.** Virtual 4C plot from fetal liver HiC data shows the interaction between *Dcbld2* promoter and *Cpox* 3’ region. *Dcbld2* promoter as anchor. **F.** 3C-qPCR result shows the interaction of TAD boundaries after *Cpox* knock down. *Dcbld2* 3’end region (Left TAD boundary) as anchor. TAD boundary regions are highlighted in green. Three biological replicates for shCPOX-4, four biological replicates for shCTRL and shCPOX-5. Data are mean ± s.d.. *: P ≤ 0.05; **: P ≤ 0.01, ***: P ≤ 0.001, ****: P ≤ 0.0001, unpaired two-tailed t-test.

Next, we checked the intra-super enhancer interactions. *CpoxeRNA* shows the highest interaction with the *Cpox* gene body 2 region and this interaction decreased after *Cpox* mRNA KD (Fig. 2B). These results demonstrate that knocking down *Cpox* mRNA can promote release of *St3gal6* and *CpoxeRNA* from *Cpox* gene body interactions, which correlates with transcription up-regulation, supporting that the gene body region of *Cpox* functions as a repressor.

### *Cpox* mRNA knock down alters interaction between TAD boundary genes

The *Dcbld2* gene is located at the left TAD boundary, with the 3’ region overlapping the TAD boundary, which contains tandem CTCF binding sites (Supplementary Fig. 2D). The *Cpox* gene is located at the right TAD boundary, with the 5’ region overlapping the boundary, but without a CTCF binding site (Supplementary Fig. 2D). Since a promoter can act as the anchor of a TAD(10,47), it is possible that the right TAD boundary is anchored by the promoter of *Cpox*. To investigate *Dcbld2*/left TAD boundary long range interactions, we first set the anchor for 3C at the *Dcbld2* promoter and found that the *Dcbld2* promoter has a higher interaction frequency with the gene body region of *Cpox* than with the *Cpox* promoter and termination site, especially with gene body region 2 (Fig. 2C). Gene body region 2 contains the 3C fragment that extends from the end of exon 4 and includes most of the last (7^th^) exon. This pattern is confirmed by virtual 4C analysis of cell cycle in situ HiC data in G1ER undifferentiated erythroid cells and in situ HiC in fetal liver erythroid progenitor cells(34,35,40) (Fig. 2D, E). After knocking down *Cpox* mRNA, the interaction between the *Dcbld2* promoter and the *Cpox* gene body region 1 and 2 decreased, whereas the interaction of *Dcbld2* promoter with the left TAD boundary significantly increased (Fig. 2C). This left TAD boundary 3C fragment overlaps with the 3’end of the *Dcbld2* gene. These results suggest that the *Dcbld2* promoter was sequestered by the gene body region of *Cpox*, and after *Cpox* mRNA loss, the *Dcbld2* promoter is released and has increased interaction with its own transcription termination site (TTS). Furthermore, the promoter interaction profiles of *Dcbld2*, *St3gal6* and *CpoxeRNA* suggest that the mechanism underlying their activation after *Cpox* KD differs from the Enhancer Release and Retargeting (ERR) model.

To investigate TAD organization in more detail, we used the left TAD boundary as an anchor for a 3C experiment. Our analysis revealed that the left TAD boundary region had the highest interaction frequency with the *Cpox* gene body region 1, which overlaps with the right TAD boundary, and this interaction was significantly increased after *Cpox* mRNA knockdown (Fig. 2F). These findings suggest that loss of *Cpox* mRNA can reinforce TAD corner interaction. Moreover, the *Cpox* gene body region 1, located at the right TAD boundary, overlaps with H3K27ac marks. The increased interaction with the left TAD boundary (*Dcbld2* 3’ region) brings the *Dcbld2* 3’ region to a conducive environment for transcription. Simultaneously, the increased interaction of the *Dcbld2* 3’ region with its own promoter positions the *Dcbld2*-1 promoter closer to this positive environment, potentially facilitating the transcription of *Dcbld2* long isoform (*Dcbld2-1*). Taken together, these results show that the TAD anchor genes *Dcbld2* and *Cpox* are interconnected head to tail. TAD anchors are typically organized with convergent CTCF sites. However, the conformation of TAD anchors that are not occupied by CTCF is not well understood. In this study, we present an example of a TAD anchored by two genes that lie in the same direction along the linear genome and are organized in a head-to-tail inter-gene connection to anchor the TAD.

### Increased transcriptional R loop formation strengthens TAD boundary insulation

The R loop has been reported to play a role in mediating TAD boundary interactions(14). To investigate whether R loop formation mediates the increased TAD boundary interaction observed after *Cpox* mRNA knockdown, we utilized a R loop-specific antibody, S9.6, and found that R loop formation was increased at both TAD boundaries after *Cpox* knockdown (Supplementary Fig. 3A). The increase in R loop formation in *Dcbld2* correlated with up-regulation of its transcription. This suggests that transcriptional activation may mediate the formation of transcriptional R loops, ultimately leading to the strengthening of the TAD corner interaction.

To further confirm the role of increased transcription in strengthening the TAD boundary interaction at the *Dcbld2* locus, we examined 4DN HiC data from other cell types and observed that CH12.LX cells, which also have this TAD, exhibit lower corner loop interaction compared to G1ER erythroid cells(34) (Supplementary Fig. 3B). RNA-seq data revealed lower *Dcbld2* and *Cpox* expression levels in CH12.LX cells compared to G1ER cells (Supplementary Fig. 3C). These results support the notion that increased TAD boundary transcription correlates with increased TAD corner interaction.

### PRC2 translocation from mRNA to chromatin to mediate the activation of neighboring genes

Next, we investigated the mechanism underlying the activation of neighboring genes and *CpoxeRNA* after *Cpox* mRNA knockdown. It has been reported that mRNA competes with chromatin to bind PRC2, and mRNA loss results in the accumulation of H3K27me3 in chromatin (4,5). To test whether PRC2 is involved in the activation of neighboring genes and *CpoxeRNA*, we first checked if PRC2 is bound to *Cpox* mRNA by using published dCLiP-seq data from mouse embryonic stem cells(48) and found *Cpox* mRNA interacts with PRC2 components EZH2 and SUZ12 (Fig. 3A). This interaction is confirmed by fCLiP-qPCR using antibodies against PRC2 components EZH2 and SUZ12 in iMEL cell, and we found loss of binding of PRC2 with *Cpox* mRNA in the *Cpox* knock down samples (Fig. 3B). Next, we used ChIP-qPCR to detect changes in the H3K27me3 levels at the *Cpox* locus after *Cpox* mRNA knock down (see supplementary Fig. 4A for primer locations). As shown in Fig. 3C, H3K27me3 accumulated at the *Cpox* genomic locus after *Cpox* mRNA KD, with especially significant increases at *Cpox* promoter and intron 5 regions. We then performed ChIP-qPCR to detect EZH2, the catalytic component of the PRC2 complex, and found that it also tended to be higher across *Cpox* after mRNA loss (Fig. 3D). These results show that after *Cpox* mRNA loss, PRC2 translocate from mRNA to the *Cpox* genomic locus and this correlates with acquisition of the repressive H3K27me3 mark across *Cpox*.

**Figure 3.**
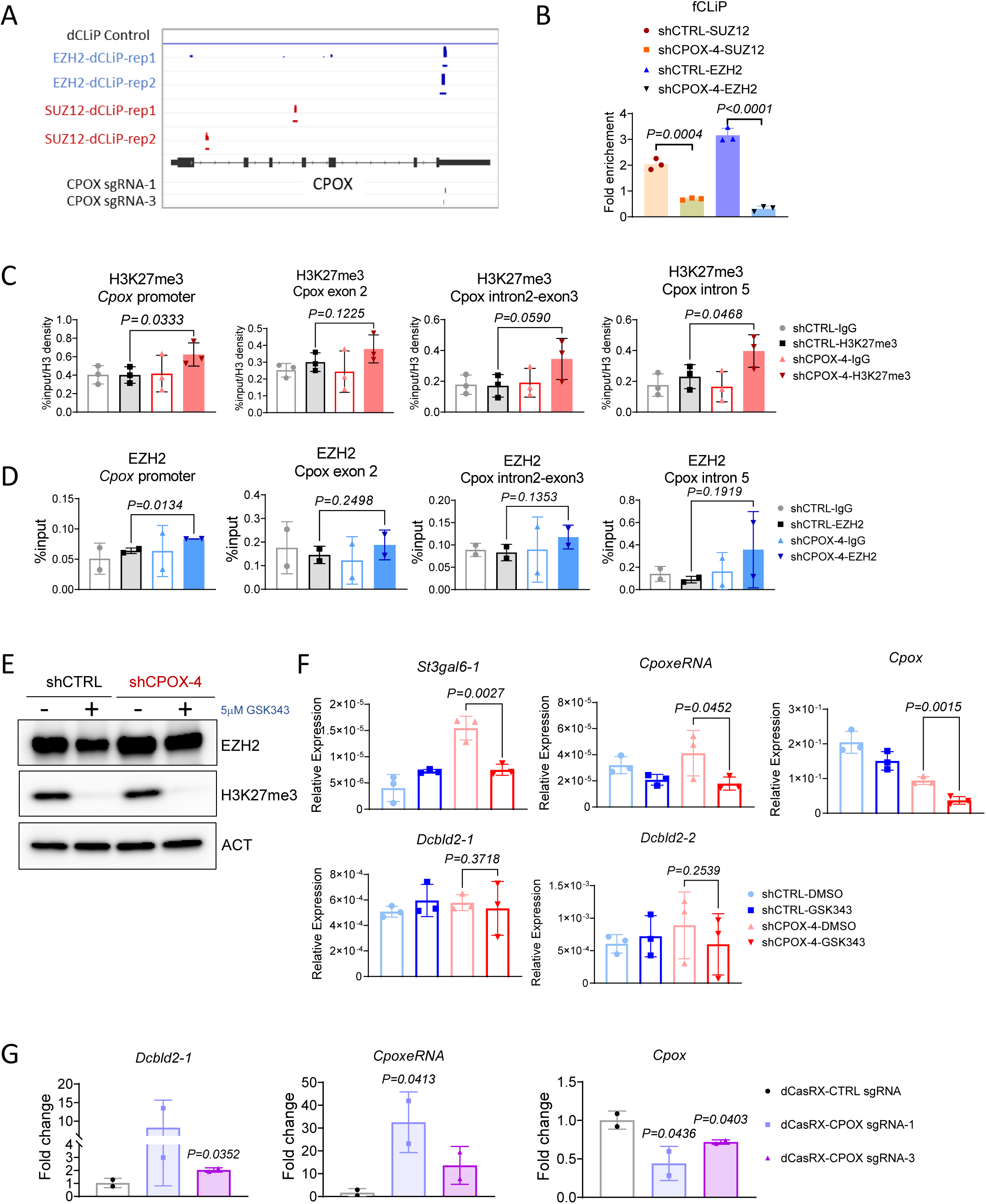
H3K27me3 and PRC2 translocation after loss of *Cpox* mRNA. A. IGV tracks show EZH2 and SUZ12 dCLIP result and location of sgRNAs used in dCasRX experiment. **B.** fCLiP-qPCR result in iMEL shows the binding of PRC2 components to *Cpox* mRNA. n = 3 technical replicates. Data are mean ± s.d., unpaired two-tailed t-test. **C.** ChIP-qPCR result shows H3K27me3 accumulation at the *Cpox* locus after *Cpox* knock down in iMEL cells. Three biological replicates. Data are mean ± s.d., unpaired one-tailed t-test. **D.** ChIP-qPCR result shows EZH2 accumulation at the *Cpox* promoter after *Cpox* knock down in iMEL cells. Three biological replicates. Data are mean ± s.d., unpaired one-tailed t-test. **E.** Western blot result shows depletion of H3K27me3 after 24h GSK343 treatment. **F**. qRT-PCR result shows gene expression change after 24h GSK343 treatment in UMEL. Vehicle is treated with DMSO only. Statistical test was performed with DMSO-shCPOX-4 and GSK343-shCPOX-4 samples. Three biological replicates. Data are mean ± s.d.; unpaired one-tailed t-test. **G.** qRT-PCR result shows gene expression level in control and dCasRX-CPOX UMEL cells. Two biological replicates, data are mean ± s.d., one-tailed t-test.

To further confirm the activation of neighboring genes is due to PRC2 mediated repression of the *Cpox* genomic locus, we treated the control and *Cpox* knockdown cells with PRC2 inhibitor GSK343. Western blot analysis shows that PRC2 component EZH2 levels did not change in GSK343 treated cells, while H3K27me3 levels were nearly completely depleted (Fig. 3E), consistent with the property of this drug to inhibit the enzyme activity of EZH2 but not its protein level. After GSK343 treatment, the levels of *CpoxeRNA* and neighboring gene *St3gal6* dropped to control levels, *Dcbld2-2* also exhibits a decreased trend, confirming that their up-regulation after *Cpox* mRNA knockdown is due to PRC2 translocation and repression of the *Cpox* genomic locus (Fig. 3F).

PRC2 bound by nascent RNA was reported act *in cis* to repress the expression of neighboring genes(48). From published dCLiP-seq data(48), SUZ12 binds at the first and third introns of *Cpox* nascent RNA, while EZH2 binds at the last exon, which is within the 3C fragment region (*Cpox* gene body 2) that has decreased interaction with *CpoxeRNA* and *Dcbld2* after *Cpox* mRNA knockdown (Fig. 3A). This suggests that EZH2 bound at *Cpox* mRNA might have a role in inhibiting *CpoxeRNA* and *Dcbld2*, likely due to the close spatial proximity of their genomic loci. Moreover, *Cpox* mRNA may act as a bridge to facilitate the recruitment of EZH2 to the *CpoxeRNA* and *Dcbld2* loci. To further confirm this hypothesis, we used dCasRX with two different sgRNAs targeting the EZH2 binding site on *Cpox* mRNA, thereby preventing its interaction with *Cpox* mRNA. As shown in Fig. 3G, when using primers detecting *Dcbld2* long isoform, which are located near the H3K27me3 peak, we observed a significant upregulation of *Dcbld2* long isoform and *CpoxeRNA* in the dCasRX treated sample. However, *Cpox* expression decreased slightly in the dCasRX treated samples. In UMEL cells, knockdown of *Cpox* mRNA did not upregulate the long isoform of *Dcbld2* (Fig. 1C). Therefore, the upregulation of the *Dcbld2* long isoform observed in the dCasRX treated cells can be attributed to the loss of EZH2 binding to *Cpox* mRNA. These findings suggest that EZH2 binding to mRNA plays a repressive role *in trans* to inhibit the distal target genomic locus.

### Perturbed p300 binding at *Cpox* Intron 5 contributes to boundary gene *Dcbld2* and intra-TAD enhancer activation

The 3C fragment located in the 3’ region of *Cpox* interacts with both the *Dcbld2* promoter and *CpoxeRNA*. It spans from exon 4 of *Cpox* to include most of its last exon. Intron 5 of *Cpox* contains binding sites for known transcription factors (TFBS) that can regulate transcription termination(49), including GATA1, TAL1 and p300 (Fig. 4A). We reasoned that intron 5 TFBS plays an important role in regulating *Dcbld2* and *CpoxeRNA* expression. To test this hypothesis, we used CRISPR-Cas9 to delete intron 5 TFBS (supplementary Fig. 4B to D). After intron 5 TFBS deletion, the mRNA and protein levels of *Cpox* exhibited different patterns in different single cell clones, as shown in Supplementary Fig. 5A and B. This clonal variability is probably due to heterogeneity of the wild type MEL cell population(50). However, *Dcbld2*, *St3gal6* and *CpoxeRNA* were consistently up-regulated (Fig. 4B), indicating that intron 5 TFBS has an inhibitory role. This result argues that activation of *Dcbld2*, *St3gal6* and *CpoxeRNA* after *Cpox* mRNA knockdown is due to the loss of the intron 5 locus, and not due to an altered level of CPOX protein.

**Figure 4.**
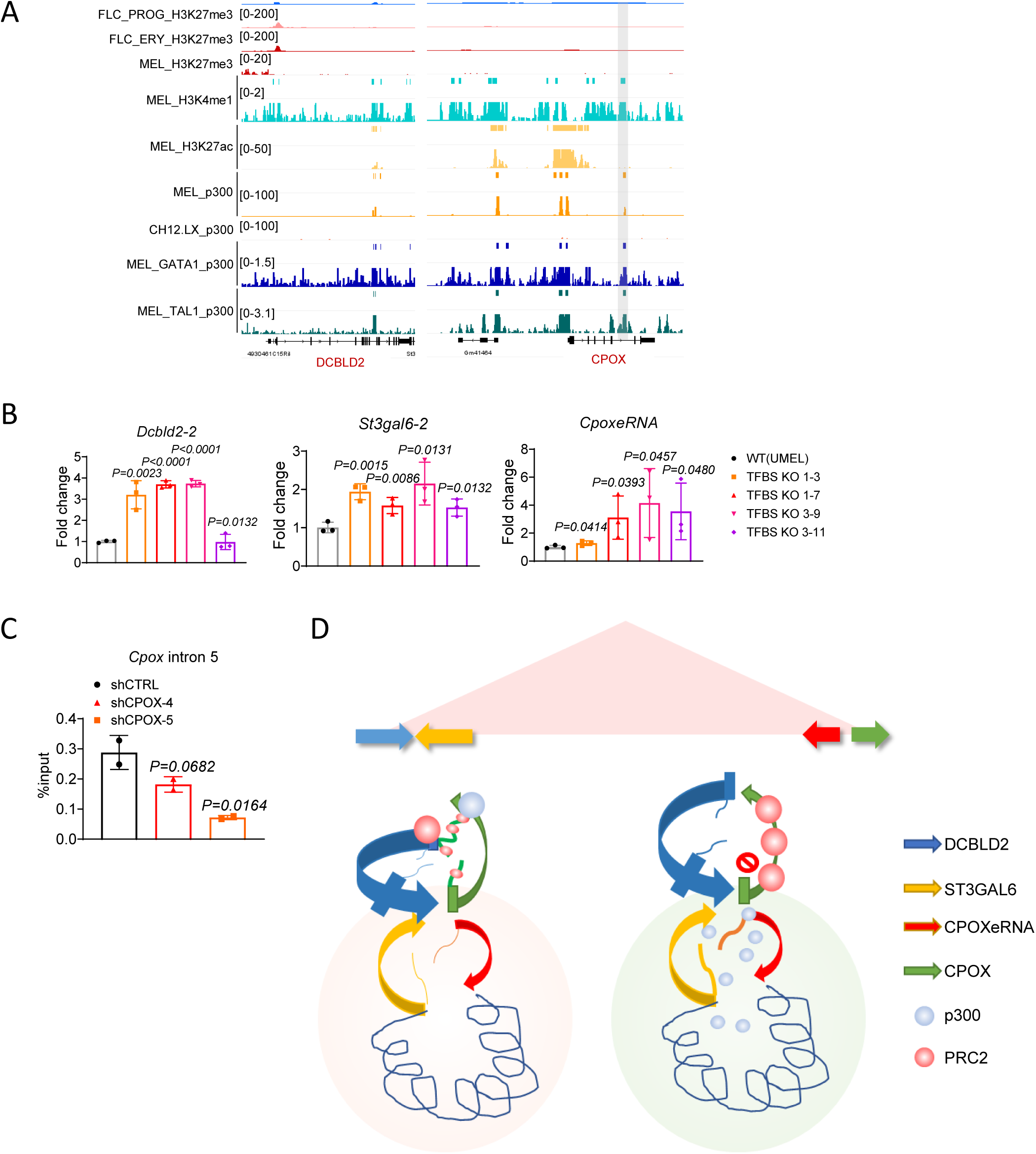
p300 binding at *Cpox* intron 5 contributes to the regulation of intra-TAD enhancer and TAD boundary gene *Dcbld2*. A. UCSC genome browser tracks shows p300, H3K27ac, H3K27me3, and H3K4me1 pattern at *Dcbld2* promoter and *Cpox* intron 5. ChIP-seq data from ENCODE. p300 only peak was highlighted in grey. **B.** qRT-PCR result shows the expression of *Dcbld2-2*, *St3gal6-2* and *CpoxeRNA* after *Cpox* intron 5 TFBS deletion in UMEL cells. Three biological replicates. Data are mean ± s.d., unpaired one-tailed t-test. **C.** ChIP-qPCR result shows p300 binding at the *Cpox* intron 5 region after *Cpox* knock down by shRNA in UMEL cells. Two biological replicates. Data are mean ± s.d., unpaired one-tailed t-test. **D.** Model of *Cpox* mRNA loss activate neighboring gene and enhancer. Left graph shows in normal situation, right graph shows *Cpox* mRNA knock down situation. Colored arrows represent the genomic locus of the protein genes and enhancer. Colored curves represent the corresponding RNA transcribed. Blue dots represent p300, orange dots represent PRC2. Rectangles on genes represent the promoters.

**Figure 5.**
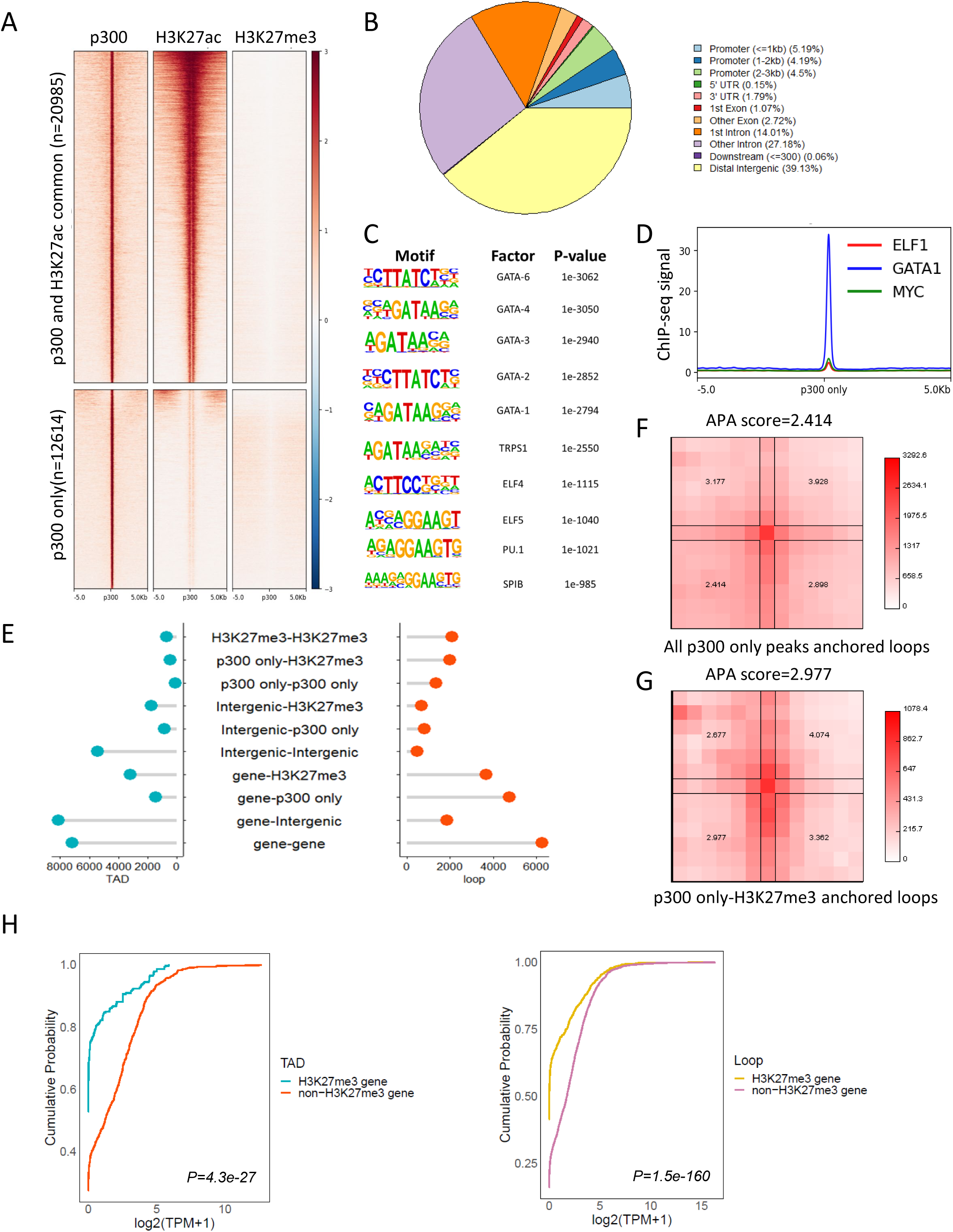
Genome wide discovery of p300-H3K27me3 repressive loop. A. Heatmap shows the distribution of p300, H3K27ac and H3K27me3 signals around +/-5kb region of p300 only peak and p300/H3K27ac common peaks. **B.** Pie chart shows the distributions of annotated features around p300 only peaks in MEL cell. **C.** Homer motif analysis identifies significant enrichment of transcription factors binding sites at p300 only peaks. **D.** Binding profiles of transcription factors at p300 only peak in MEL cells. **E.** Distribution of different types of interaction observed at TAD boundary(blue) and chromatin loops(red). X axis represents number of different types of interactions. “Gene” represents genes that do not overlap with H3K27me3 and p300 only peak. “Intergenic” represents intergenic regions that do not overlap with H3K27me3 and p300 only peak. **F** and **G.** Aggregated peak analysis (APA) of chromatin loops anchored by p300 only peaks **(F)** and p300 only peak-H3K27me3 peaks **(G)** in G1ER late G1 cell. HiC matrix was normalized with VC_SQRT, window = 6. **H.** ECDF plot shows the comparison of the expression level of H3K27me3 non-overlapping and overlapping genes looped to p300 only peaks at TAD boundary or Chromatin loops. MEL cell RNA-seq processed expression quantification data was obtained from ENCODE. Wilcoxon test was used to compare the log2(TPM+1) value of H3K27me3 genes and non-H3K27me3 genes, P values ≤ 0.05 indicating a significant difference.

Next, we investigated which protein is responsible for the repression of intra-TAD neighboring genes and enhancer. By examining the ENCODE ChIP-seq data in MEL cell, we found a p300 peak at the intron 5 TFBS. This p300 peak overlaps with H3K4me1, but not with H3K27ac (p300 only peak, Fig. 4A). Regulatory elements with H3K4me1 but without H3K27ac were described as poised enhancers. Some poised enhancers also contain H3K27me3(51-54), while other regulatory elements with H3K4me1 and H3K27me3 were considered silencers(55). However, according to our KO experiment result, the intron 5 element seems to behave like a repressor but without overlap with H3K27me3. p300 is known to catalyze and deposit H3K27ac, which is a hallmark active epigenetic mark(56,57). However, recent research has suggested that p300 can also occupy the H3K27me3 region at poised enhancers(52,58,59). Additionally, it has been reported that EZH2, a key component of the polycomb repressive complex 2 (PRC2), interacts with p300(60). Notably, p300/CBP has been found to be crucial for PRC2 silencing in regions that are co-occupied by CBP/PRC2, independent of its enzymatic function(61). However, the role of p300 occupancy in the absence of H3K27ac modifications is not fully understood. By examining the ENCODE histone ChIP-seq data, we found that the *Dcbld2* promoter is enriched in H3K27me3 in MEL cells (Fig. 4A). Given that the *Dcbld2* promoter and *Cpox* intron 5 TFBS exhibit interaction, we postulated that this association may be facilitated by PRC2-p300 and p300 bound at *Cpox* intron 5 should have a repressive role due to its association with PRC2. If this assumption is valid, we would expect that p300 binding at the *Cpox* intron 5 TFBS would diminish following *Cpox* mRNA depletion. Our findings confirm this hypothesis, as evidenced by the observed reduction in p300 binding at the *Cpox* intron 5 TFBS after *Cpox* mRNA knockdown (Fig. 4C).

In conclusion, the observed decrease in p300 binding at *Cpox* intron 5 following *Cpox* mRNA knockdown implies the release of p300 from the *Cpox* intron 5 locus (Fig. 4D). This event may subsequently lead to an increase in the concentration of activators in the insulated TAD environment, thereby activating enhancers and protein coding genes located in the vicinity. P300 release may also explain the activation of intra-TAD genes (*Dcbld2-2* and *St3gal6*) and enhancer after CPOX TSS deletion.

### Genome-wide discovery of p300-H3K27me3 loops unveils their repressive role in regulating gene expression

Next, we sought to determine the prevalence of p300-PRC2/H3K27me3 loops across the genome. We analyzed ENCODE MEL cell ChIP-seq profiles for the genome-wide distribution of H3K27me3 and extended H3K27ac (+/- 1kb region) peaks around p300 binding sites. We found that a subset of p300 binding sites did not exhibit occupancy by H3K27ac (Fig. 5A). Notably, both the H3K27ac overlapping peak and the H3K27ac independent peak were devoid of H3K27me3 binding. To further explore this phenomenon, we focused on p300 peaks that did not overlap with the extended H3K27ac peaks (+/- 1kb region) for further analysis. These H3K27ac-independent p300 peaks were predominantly localized in intergenic and intronic regions, with a notable preference for other introns besides the first intron (Fig. 5B). Given that p300 has diverse targets beyond histone H3K27ac, we performed motif analysis using HOMER to identify the transcription factors co-localized at p300-only peaks. Our analysis revealed a significant enrichment of cell type-specific transcription factors, including, primarily GATA factors, but also ELF factors and PU.1 (Fig. 5C). To validate these findings, we examined the binding profiles of several transcription factors, confirming the enrichment of cell type-specific transcription factors at the p300-only peaks (Fig. 5D).

Furthermore, we aimed to identify the loops anchored by both p300 and PRC2/H3K27me3 on a genome-wide scale. To accomplish this, we analyzed published G1ER cell cycle Hi-C data (34,40). we found that 1.52% of TAD boundary interactions and 8.33% of loops were anchored by p300- only and H3K27me3 peaks (Fig. 5E). The formation of loops anchored by all the p300-only peaks and p300-only-H3K27me3 peaks was confirmed through aggregated peak analysis (APA) (Fig. 5F and G).

To further investigate whether p300-only-H3K27me3 anchored loops have a repressive role, we compared the expression levels of genes that form loops with p300-only peaks. Notably, genes overlapping with H3K27me3 exhibited significantly lower expression levels compared to those without H3K27me3 peaks (Fig. 5H). This finding confirms that p300-H3K27me3/PRC2 loops are associated with gene repression. Thus, we suggest that p300-H3K27me3 loops could serve as a mechanism for protein-coding genes to repress their interacting protein-coding counterparts, providing insight into the regulatory mechanisms underlying gene expression.

### Knock down of non-oncogenes in a head-to-tail TAD boundary gene pair activates its partner oncogene

Next, we sought to explore whether the head-to-tail conformation of boundary genes regulation is pervasive and whether it could exist beyond super enhancers. Since the TAD is a conserved structure, we extended our analysis to different cell types and species. We examined the interaction patterns of TAD boundary genes in human K562 cells and GM12878 cells, and mouse CH12.LX cells and G1ER cells, and we found that the most enriched interaction pattern among TAD boundary gene pairs are gene body-gene body interaction and promoter-gene body interaction (PG) (Fig. 6A and Supplementary Fig. 6A). When comparing the expression of promoter-gene body (PG), promoter-terminator (PT) and promoter-promoter (PP) gene pairs, genes with their gene body or terminators overlapping with a TAD boundary were expressed at a lower level than genes having their promoters overlapping with TAD boundary (Fig. 6B and Supplementary Fig. 6B). This is exemplified by the *Dcbld2*-*Cpox* pair: the *Dcbld2* gene body overlaps with the TAD boundary and is expressed at a lower level than *Cpox*, whose promoter overlaps the partner boundary. This indicates the head-to-tail inter-gene interaction functions to restrict gene expression. Next, we explored whether the head-to-tail inter-gene interaction plays any role to repress oncogene activation. One example oncogene/non-oncogene pair is shown in Figure 6C, *CDK12*-*PSMD3*, which has the same conformation as *Dcbld2*-*Cpox* pair with gene body of *CDK12* and promoter of *PSMD3* overlap with TAD boundary. *CDK12* is an oncogene which is up-regulated and has highest expression level in Acute Myeloid Leukemia (AML) (Figure 6D), while its partner boundary non-oncogene *PSMD3* shows down-regulation in AML patients (Figure 6E). To further validate whether there is a causal relationship between decreased expression of *PSMD3* and increased expression of *CDK12*, we performed a *PSMD3* knock down experiment in 293T cell (Figure 6F), and indeed *CDK12* was activated after *PSMD3* KD. Among the non-oncogene/oncogene pairs with promoter of non-oncogene and gene body/terminator of oncogene overlapping with a TAD boundary, we selected another 3 pairs and performed knock down experiments with shRNA in 293T cell. As shown in Fig. 6F, after knocking down the non-oncogenes, only one showed up-regulation of its oncogene partner but without statistical significance. These non-oncogene/oncogene pairs are not in the same conformation as *Dcbld2*/*Cpox* pair (in the same direction on the linear genome while upstream gene has 3’ region overlap with TAD boundary and downstream gene has promoter overlap with TAD boundary) (Supplementary Fig. 7A). This suggests that head-to-tail boundary gene conformation is required to suppress an upstream oncogene by a downstream non-oncogene. By focusing on only the conformation like *Dcbld2*/*Cpox*, we identified 358 TADs in K562 cell that are in such conformation. 2.3% of the oncogenes have their gene body/terminator overlapped with the TAD boundaries and are potentially repressed by their downstream partner non-oncogenes, which have promoter overlap with the TAD boundary. We checked the mutation distribution at promoters of TAD boundary non-oncogenes by analyzing data from the International Cancer Genome Consortium (ICGC). Oncoplots showed that in 1% of about 6000 cancer samples, the non-oncogenes have mutations at their promoters (Supplementary Fig. 7B). These results suggest that head-to-tail inter-gene interaction could be a mechanism for cells to restrain expression of a subset of oncogenes.

**Figure 6.**
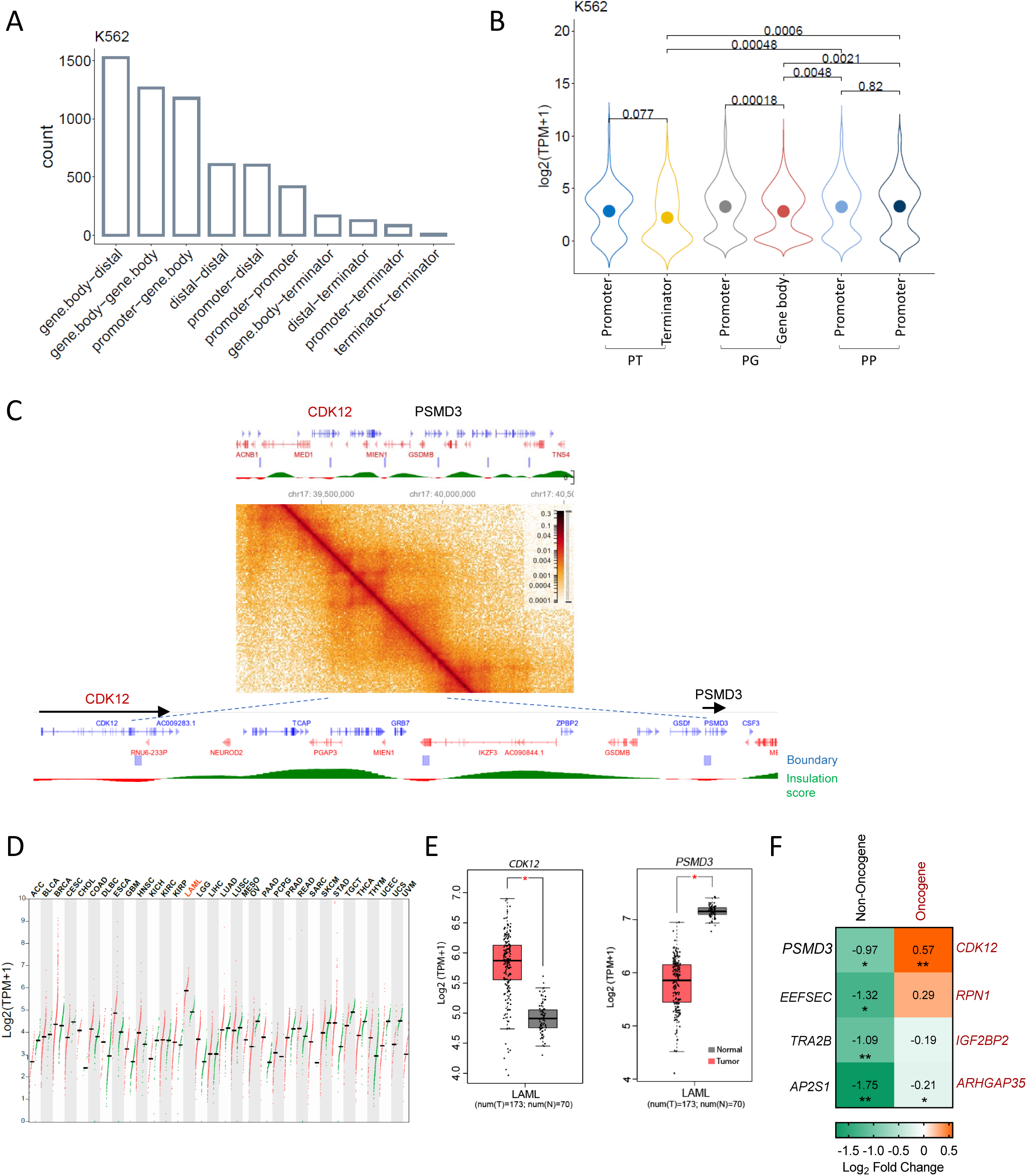
Head-to-Tail TAD boundary genes conformation is a potential mechanism to restrain the activation of certain oncogenes. A. Distribution of different types of interactions observed at TAD boundary in K562 cell. **B.** Expression level at each TAD boundary for Promoter-Terminator (PT), Promoter-Genebody (PG) and Promoter-Promoter (PP) boundary pairs in K562 cells. Dots inside of the violin plots show the mean value. Kruskal-Wallis test, P values ≤ 0.05 indicating a significant difference. C. *CDK12*-*PSMD3* TAD in K562 cell. Data from 4DN portal(8,34). **D.** Dot plot shows the expression level of *CDK12* across various tumor samples and paired normal tissues. Data from GEPIA(71). AML is highlighted in red. **E.** Expression level of *CDK12* and *PSMD3* in AML samples and paired normal tissues. Data from GEPIA(71) with Log2FC cutoff as 0.5, p value cutoff as 0.05. **F.** Expression level of TAD anchor oncogenes after knocking down its paired TAD anchor non-oncogene genes in 293T cell. Two biological replicates were shown here. *: P ≤ 0.05; **: P ≤ 0.01, ***: P ≤ 0.001, ****: P ≤ 0.0001, unpaired one-tailed t-test.

## DISCUSSION

Our findings demonstrate a novel non-coding function of mRNA, revealing its regulatory role in neighboring gene expression and chromatin structure. As illustrated in the model in Figure 4D, *Cpox* mRNA loss results in the translocation of the PRC2 complex from the mRNA to chromatin, leading to an accumulation of H3K27me3 at the *Cpox* genomic locus, thereby altering its chromatin looping pattern with neighboring genes. On one hand, both the TSS and TTS region of TAD boundary gene *Dcbld2* interact with *Cpox*, connected in an inter-gene TSS-TTS loop conformation. Following *Cpox* mRNA depletion, the *Dcbld2* TSS-*Cpox* TTS loop decreased while the *Dcbld2* TSS- *Dcbld2*-TTS loop increased. This release of the *Dcbld2* TSS from the *Cpox* TTS allows it to connect with its own TTS, thus facilitating its transcription. Furthermore, the increased transcription leads to the formation of transcriptional R loops, which, in turn, strengthen the TAD corner loop. On the other hand, p300 is released from *Cpox* intron 5 due to PRC2 translocation, resulting in an increased availability of p300 in the TAD insulated environment. Elevated p300 availability correlates with the activation of enhancer and protein coding genes within the TAD. In summary, our results reveal that mRNA plays a crucial role in gene regulation, particularly in chromatin looping patterns and enhancer activation. These findings shed light on a previously unknown mechanism of gene regulation and highlight the importance of understanding the complex interplay between mRNA and chromatin structure.

Among the individual enhancers making up a super enhance there is a hierarchy in that these elements may have variable enhancer activity(62-64). For example, super enhancer components can be divided into hub enhancers and non-hub enhancers, with hub enhancers associated with higher levels of cohesin and CTCF binding(62). For protein coding gene-containing super enhancers, the contribution to enhancer activity of a protein coding gene versus a non-coding enhancer element remains unexplored. Our results demonstrating that in the super enhancer containing the protein coding gene *Cpox* and non-coding *CpoxeRNA*, the protein coding gene seems to have a dominant role over the non-coding gene. Specifically, the protein coding gene *Cpox* represses the non-coding enhancer. Many studies have focused on how enhancers could regulate gene expression, while less attention has been paid to how the enhancer could be regulated. Our results provide an example showing that an enhancer can be regulated and harnessed by a protein coding gene and the mRNA it encodes. The ERR model shows that activation of an enhancer after promoter repression is due to loss of interaction with the promoter(3). However, in our case, we observed *CpoxeRNA* mainly interacts with the gene body region of CPOX and the interaction between *CpoxeRNA* and the *Cpox* promoter is unchanged following *Cpox* mRNA loss. Within the confines of the TAD we studied, *Cpox* mRNA loss could result in locally increased free PolII or another transcription activator, which will then have an increased chance to bind with the *CpoxeRNA* promoter and drive its expression. Our results also indicate a decrease in p300 at the *Cpox* intron 5 locus, with an increased tendency to bind at the *CpoxeRNA* locus after *Cpox* mRNA knockdown, suggesting that p300 could be a potential activator of *CpoxeRNA* after *Cpox* mRNA knock down. In addition, *Cpox* intron 5 p300 forms a chromatin loop with H3K27me3/PRC2 at the *Dcbld2* promoter. This may indicate p300 at *Cpox* intron 5 has a repressive role to harness *CpoxeRNA* expression.

We propose two plausible mechanisms for repression of *Dcbld2* by *Cpox*, (1) sequestration of the *Dcbld2* promoter by the transcription termination region of *Cpox* and (2) a novel type of repressive chromatin loop established by p300 and PRC2/H3K27me3. These two mechanisms are not mutually exclusive, and they could exist concurrently. The two anchors of a chromatin loop are likely not in direct contact with each other but are instead spatially proximate. This spatial proximity could allow space for multi-component protein complexes, and multiple proteins that exist at the same time within the spatial region around anchors(65). PolII termination related factors might create a repressive environment, which inhibits the firing of the promoter of an interacting gene. For instance, we observed gene pairs like *PSMD3*-*CDK12* exist in TSS-TTS interaction conformation but do not possess a p300 binding site (Fig. 6). This implies that the transcription termination region/environment is sufficient to repress a promoter. With regard to the new p300-PRC2/H3K27me3 loop at the *Dcbld2*-*Cpox* pair, p300-PRC2/H3K27me3 could act additively to repress the *Dcbld2* long isoform promoter. For the discrepancy in *Dcbld2*-1 and *Dcbld2*-2 expression between UMEL and iMEL cells after *Cpox* KD, we believe that the failed induction of *Dcbld2*-1 in UMEL cells may be due to promoter competition between the two *Dcbld2* promoters. In UMEL cells, both isoforms are expressed at a similar level. The transcription activators may be sequestered and blocked by the promoter of the short isoform, as it overlaps with the TAD boundary and has shorter linear and spatial distance to positive transcription elements such as its own TTS and the enhancer (right TAD boundary), thus leading to the failed activation of *Dcbld2* long isoform after *Cpox* KD and after deletion of intron 5 TFBS. Meanwhile, in iMEL cells, the promoter of the long isoform becomes dominant, and the promoter of the short isoform is nearly inactivated. The dominant long isoform promoter can now access the positive transcription environment after *Cpox* KD, resulting in the upregulation of the long isoform.

We uncovered a new type of loop anchored by H3K27ac-independent p300 and PRC2/H3K27me3. Although the number of this type of loop is less than 10% in the genome, it confers cell type specific chromatin looping and regulation of gene expression, since p300 only peaks and PRC2/H3K27me3 are variable in different cells. Our results suggest that this loop could be mediated by RNA, as depletion of *Cpox* mRNA reduces the frequency of this chromatin interaction. RNAi can induce nascent transcript degradation and premature transcription termination(66), and antisense oligonucleotide (ASO) mediated knock down was also reported to trigger nascent RNA degradation and premature transcription termination(67,68). We observed *Cpox* nascent RNA level decreased after shRNA knock down (Supplementary Fig. 2C) and increased R loop formation at the *Cpox* locus (Supplementary Fig. 3A), consistent with premature transcription termination occurs. Further experiments blocking the binding of EZH2 to *Cpox* mRNA showed up-regulation of *Dcbld2* (Fig. 3G). This shows that the effect we observed in knockdown experiments is due to loss of *Cpox* mRNA rather than at the level of transcription perturbation and suggests that *Cpox* mRNA binds to PRC2, which may be the source of PRC2 at the *Dcbld2* promoter. In addition, CBP/p300 was also reported to bind with RNA(69), indicating that p300 may bind to nascent *Cpox* mRNA. Therefore, we propose that *Cpox* mRNA may act as a bridge to facilitate the formation of the loop between the *Dcbld2* promoter and the *Cpox* gene body region. Furthermore, the spatial proximity of p300 and PRC2 may allow p300 to stabilize PRC2 at the *Dcbld2* promoter.

Our findings demonstrate that PRC2 plays a crucial role in mediating the non-coding function of mRNA to regulate neighboring gene expression. This highlights the possibility that mutations in mRNA that do not affect its translated protein but disrupt its ability to bind PRC2 may lead to dysregulation of neighboring genes and enhancers. Moreover, we observed that p300 only peaks bind to intronic sites, suggesting that variants at these sites could have disease relevance since they may cause aberrant expression of disease-associated genes with which they interact. These findings provide a potential mechanism for how mutations identified through genome-wide association studies (GWAS) can affect gene regulation through RNA mutations. We further show that Head-to-Tail inter-gene interaction could be clinically important. As knockdown of TAD anchor non-oncogenes led to activation of its partner boundary oncogene, RNA mutation of the non-oncogene could be a potential mechanism that underlies cancer driver gene activation. Chromatin 3D conformation alteration in cancers may create new TAD boundaries that overlap with oncogenes or tumor suppressor genes to activate or suppress their expression. We also found around 2% of tumor suppressor genes are located in TAD boundaries that connected to downstream partner boundary genes in head to tail conformation. We speculate these tumor suppressor genes could be re-activated by knock down of its partner TAD boundary gene in cancer samples, which could be a RNAi based novel therapeutic approach for cancer.

## DATA AVAILABILITY

Figure 1A Mouse fetal liver HiC data fastq files were downloaded from EBI ArrayExpress (https://www.ebi.ac.uk/arrayexpress/) accession number E-MTAB-2414.

Supplementary Fig. S1G mouse fetal liver progenitor and differentiated HiC process data was downloaded from GSE184974.

4DN .hic file used in this study are downloaded from 4DN data portal: G1ER in situ cell cycle HiC late G1 phase (4DNFI6H926RO), CH12.LX (4DNFI8KBXYNL), and K562 in situ HiC (4DNFI18UHVRO). Cell cycle G1ER in situ HiC boundary files and insulation score files are downloaded from 4DN data portal(34).

TAD files and loop files for G1ER cell were downloaded from GSE129997. TAD files for K562 cell, GM12878 cells and CH12.LX cells were downloaded from GSE63525.

RNA-seq gene quantification files were downloaded from ENCODE: MEL cell (ENCFF199TJO.tsv), K562 cell (ENCFF421TJX.tsv), GM12878 cell (ENCFF345SHY.tsv), CH12.LX cell (ENCFF164NIW.tsv).

ChIP-seq data for MEL cell H3K27ac (ENCFF972YIT.bed.gz and ENCFF847MJA.bigWig), p300 (ENCFF185UAQ.bed.gz and ENCFF116DCE.bigWig), H3K27me3(ENCFF336CRJ.bigWig), H3K9me3 (ENCFF711NWS.bigWig), ELF1 (ENCFF688GOC.bigWig), GATA1 (ENCFF841DLH.bed.gz and ENCFF222HZM.bigWig), MYC (ENCFF944GOL.bigWig), TAL1(ENCFF893PQE.bed.gz and

ENCFF660OIM.bigWig) and CH12.LX EP300 (ENCFF610VZA.bigWig) were downloaded from ENCODE. Fetal liver cell ChIP-seq and RNA-seq data were downloaded from a public dataset(70), and converted to mm10. Erythroid specific enhancers were downloaded from a public dataset(45) and liftover to mm10. ChIP-seq data for asynchronous G1ER cell CTCF and RAD21 were downloaded from 4DN data portal.

dCLiP-seq data were downloaded from GSE141700.

ICGC cancer mutations data were downloaded from ICGC data portal version—release 28.

## CODE AVAILABILITY

The source code available at Github (https://github.com/Chromatin-RNA/CPOXmRNA).

## SUPPLEMENTARY DATA

### Supplementary Figures

### Supplementary Table 1

List of primers, shRNAs, sgRNAs, and LNA ASOs used in this study. This table contains the sequence of primers, shRNAs, sgRNAs, and LNA ASOs used in this study.

## ACKNOWLEDGEMENTS

We thank Dr. Vittorio Sartorelli for reviewing the manuscript and helpful suggestions, and lab member for feedback on the manuscript. The authors acknowledge the use of the computational resources of the NIH HPC Biowulf cluster (http://hpc.nih.gov).

## Author contributions

B.X. conceived the study, designed and performed all the experiments and bioinformatics analysis, and wrote the manuscript. A.D. acquired funding, supervised the study, analyzed the data and wrote the manuscript.

## Materials and correspondence

Correspondence and requests for materials should be addressed to B.X. or A.D.

## FIGURE LEGENDS

**Supplementary Figure 1.**
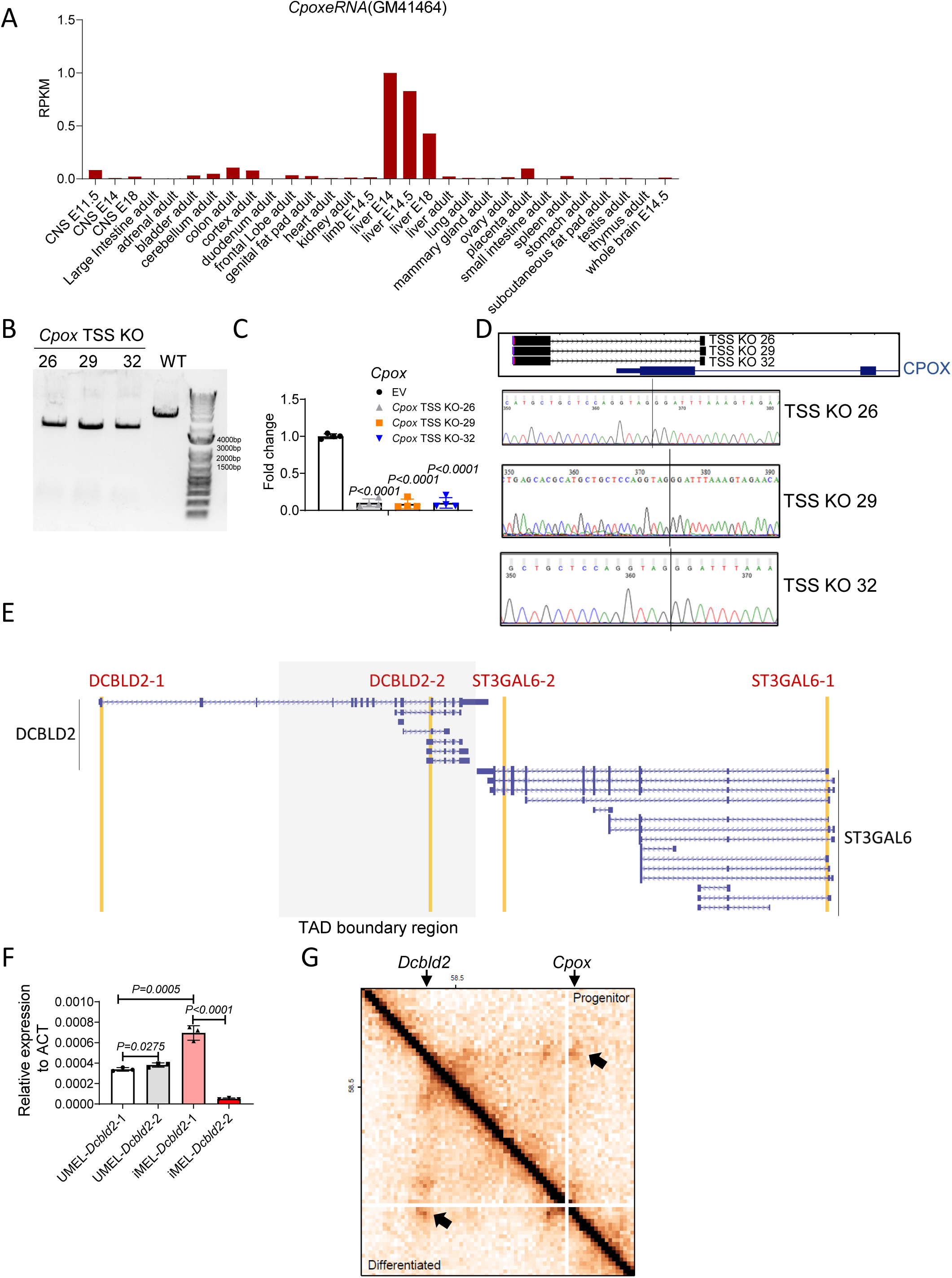
*CpoxeRNA* tissue expression, *Cpox* TSS deletion, and primer location. **A.** Tissue expression pattern of *CpoxeRNA* (*Gm41464*). Data from NCBI. **B.** Gel image for *Cpox* TSS KO confirmation. **C**. *Cpox* mRNA expression level after TSS Knock out. EV, empty vector. Data are mean ± s.d.; four biological replicates. Unpaired two-tailed t-test. **D.** Sanger sequencing result for *Cpox* TSS KO, black line shows the deletion site. **E**. Location of primers for *Dcbld2* and *St3gal6* qRT-PCR. UCSC genome browser tracks show the isoforms of *Dcbld2* and *St3gal6*. Two primer pairs each used to detect *Dcbld2* and *St3gal6* are heighted in yellow. TAD boundary region is highlighted in grey. **F.** Expression level of *Dcbld2-1* and *Dcbld2-2* in undifferentiated (UMEL) and differentiated (iMEL) MEL cells. Data are mean ± s.d.; three biological replicates. Unpaired one-tailed t-test. **G.** 5kb resolution HiC map shows the *Dcbld2-Cpox* TAD in undifferentiated and differentiated Fetal Liver cell, data from Bi et al.(39) Black arrow shows the TAD corner loop.

**Supplementary Figure 2.**
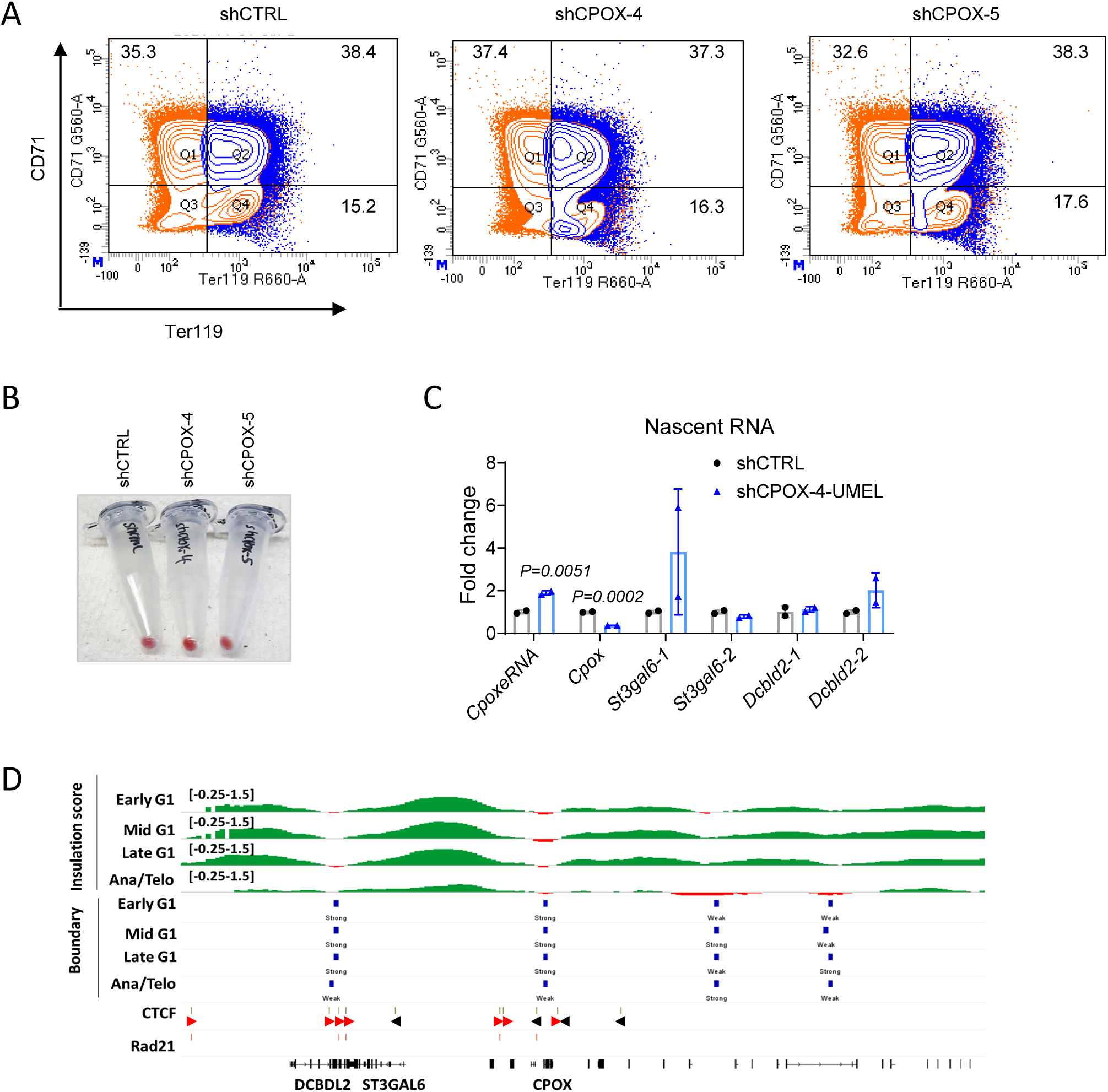
*Cpox* knock down does not affect erythropoiesis. **A.** CD71/Ter119 flow-cytometric result of differentiated shCTRL, shCPOX-4, shCPOX-5 cells. **B.** shCTRL, shCPOX-4, shCPOX-5 cells differentiated 5 days with 2% DMSO. **C.** qRT-PCR result shows nascent RNA expression level in shCTRL and shCPOX-4 UMEL cells. Two biological replicates. Data are mean ± s.d., unpaired one-tailed t-test. **D.** Insulation score and TAD boundary data from G1ER cell cycle in situ HiC experiment shows *Dcbld2* and *Cpox* overlap with TAD boundary. CTCF and RAD21 ChIP-seq data from asynchronized G1ER cell from the same study as G1ER cell cycle in situ HiC experiment were shown below.

**Supplementary Figure 3.**
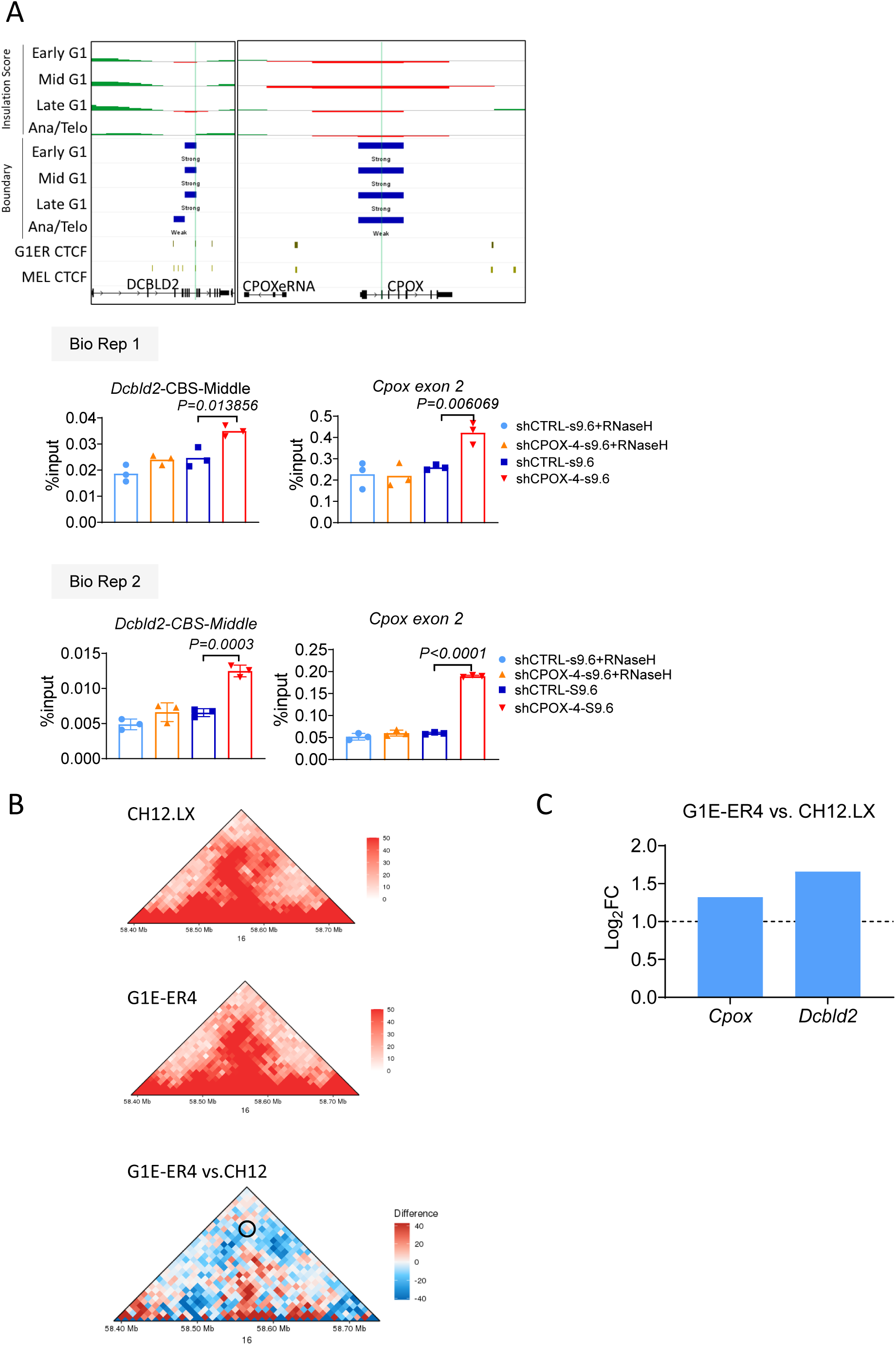
Transcription at TAD boundary correlates with TAD insulation strength. **A.**Upper panel shows the insulation score, TAD boundary, and CTCF position in G1ER and MEL cells. Lower panel shows the DRIP-qPCR result of R loop formation at the TAD boundaries after *Cpox* mRNA knock down in UMEL cells. Position of primers are highlighted in the upper panel. Data are mean ± s.d.; unpaired one-tailed t-test with three technical replicates. **B.** HiC matrix shows the *Dcbld2*-*Cpox* TAD in G1ER and CH12.LX cells. Differential matrix was shown below, TAD boundary loop was highlighted with circle in the differential matrix. **C.** Fold change of *Cpox* and *Dcbld2* expression level in G1ER and CH12.LX cell. Expression data from ENCODE RNA-seq data.

**Supplementary Figure 4.**
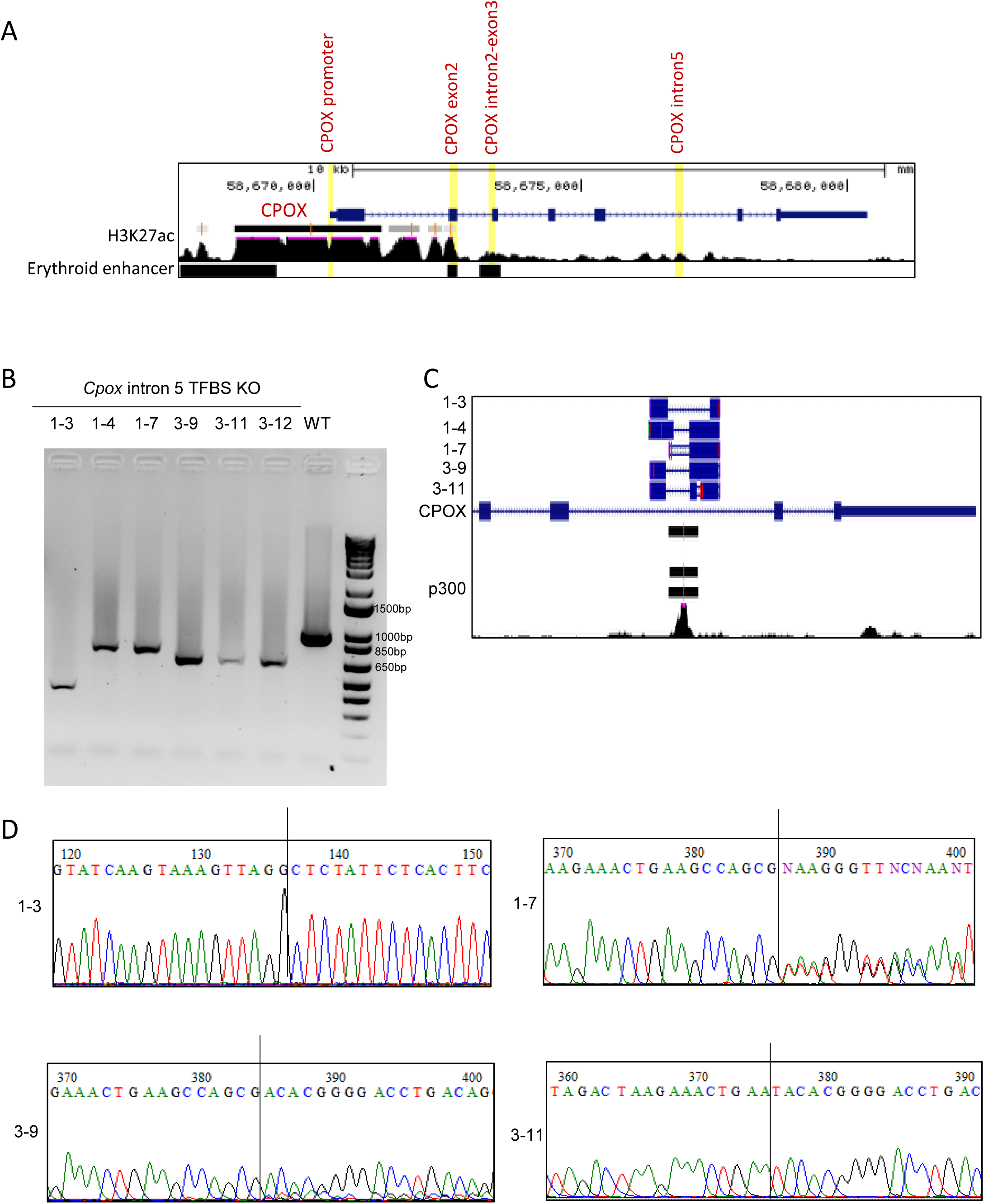
ChIP-qPCR primer location and Sanger sequencing of genetic deletion of *Cpox* intron 5 TFBS. **A.**Location of primers used for ChIP-qPCR in Figure 3 C and D. **B.** *Cpox* intron 5 TFBS KO gel image. **C.** *Cpox* intron 5 TFBS KO BLAST result, MEL p300 ChIP seq data from ENCODE. **D.** Sanger sequencing data, black line shows the deletion site.

**Supplementary Figure 5.**
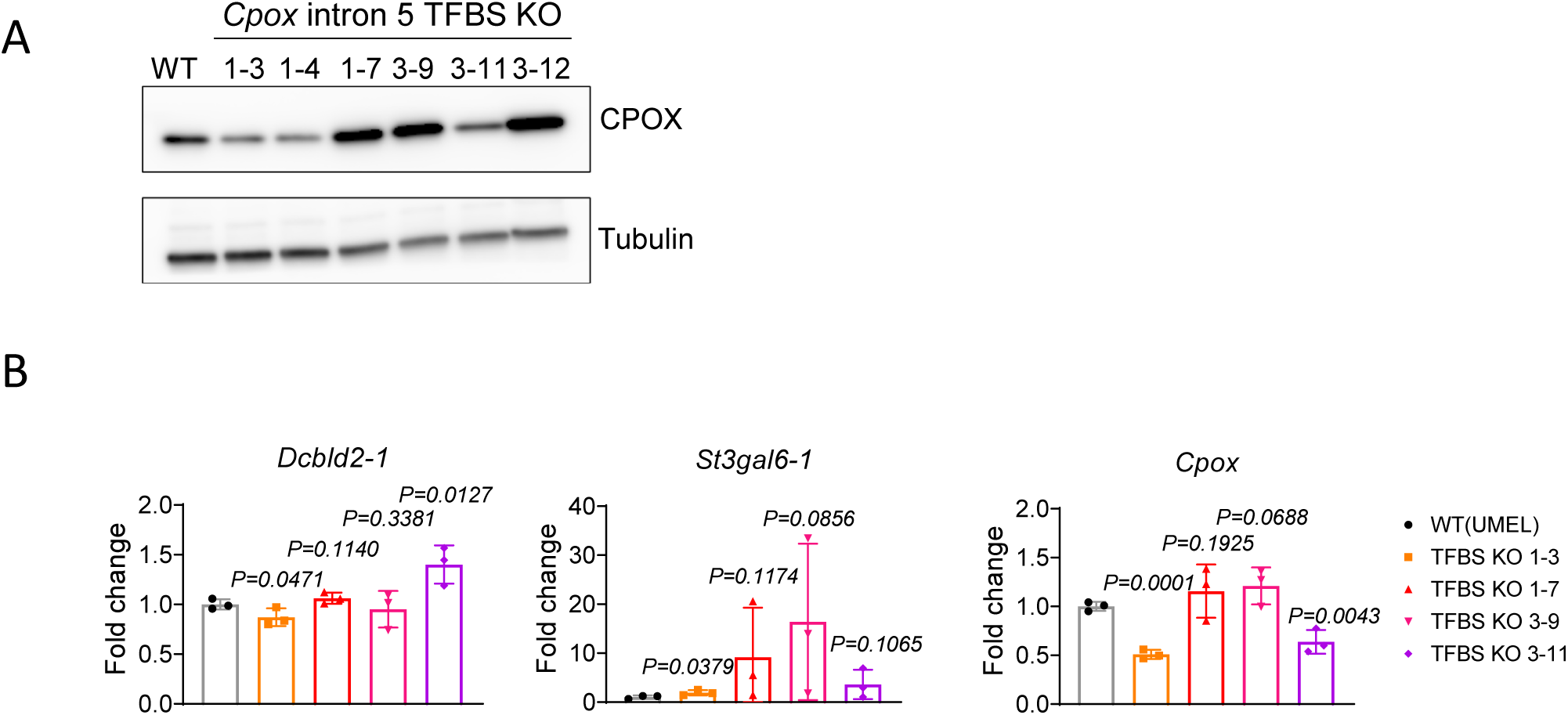
*Cpox* intron5 TFBS deletion activates intra-TAD enhancer and TAD boundary gene *Dcbld2*. **A.**Western blot result shows CPOX protein level in WT and *Cpox* intron 5 TFBS depleted cells. **B.** qRT-PCR result shows the expression of *Cpox*, *St3gal6*-1, and *Dcbld2*-1 after *Cpox* intron 5 TFBS deletion in UMEL cells. Three biological replicates. Data are mean ± s.d., unpaired one-tailed t-test.

**Supplementary Figure 6.**
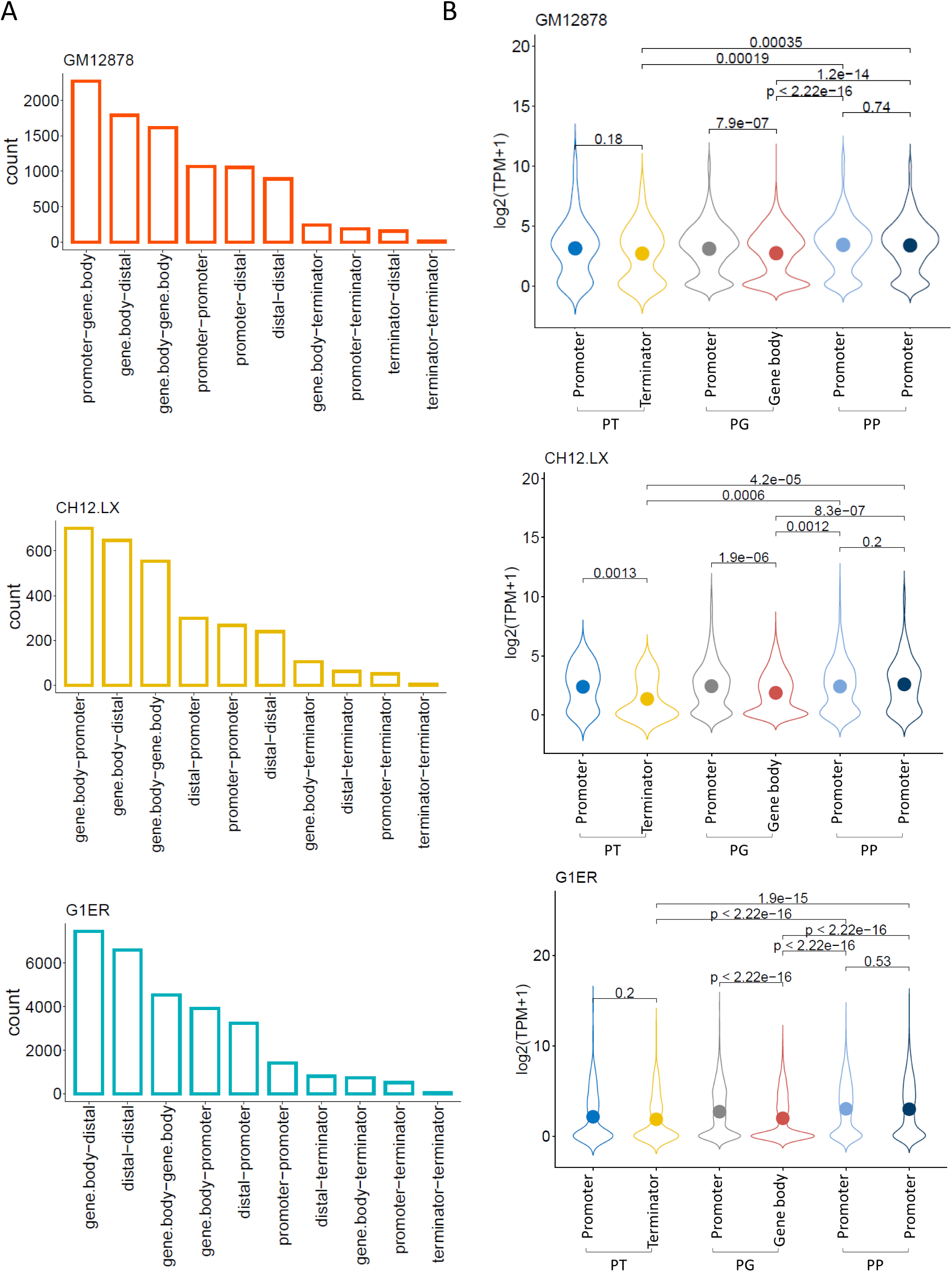
Distribution of TAD boundary gene interaction pattern and expression. A. Bar plot shows the distribution of different types of interaction observed at TAD boundary in GM12878, CH12.LX, and G1ER cells. **B.** Expression level at each TAD boundary for Promoter-Terminator (PT), Promoter-Genebody (PG) and Promoter-Promoter (PP) boundary pairs in GM12878, CH12.LX, and G1ER cells. Dots inside of the violin plots show the mean value. Kruskal-Wallis test, P values ≤ 0.05 indicating a significant difference.

**Supplementary Figure 7.**
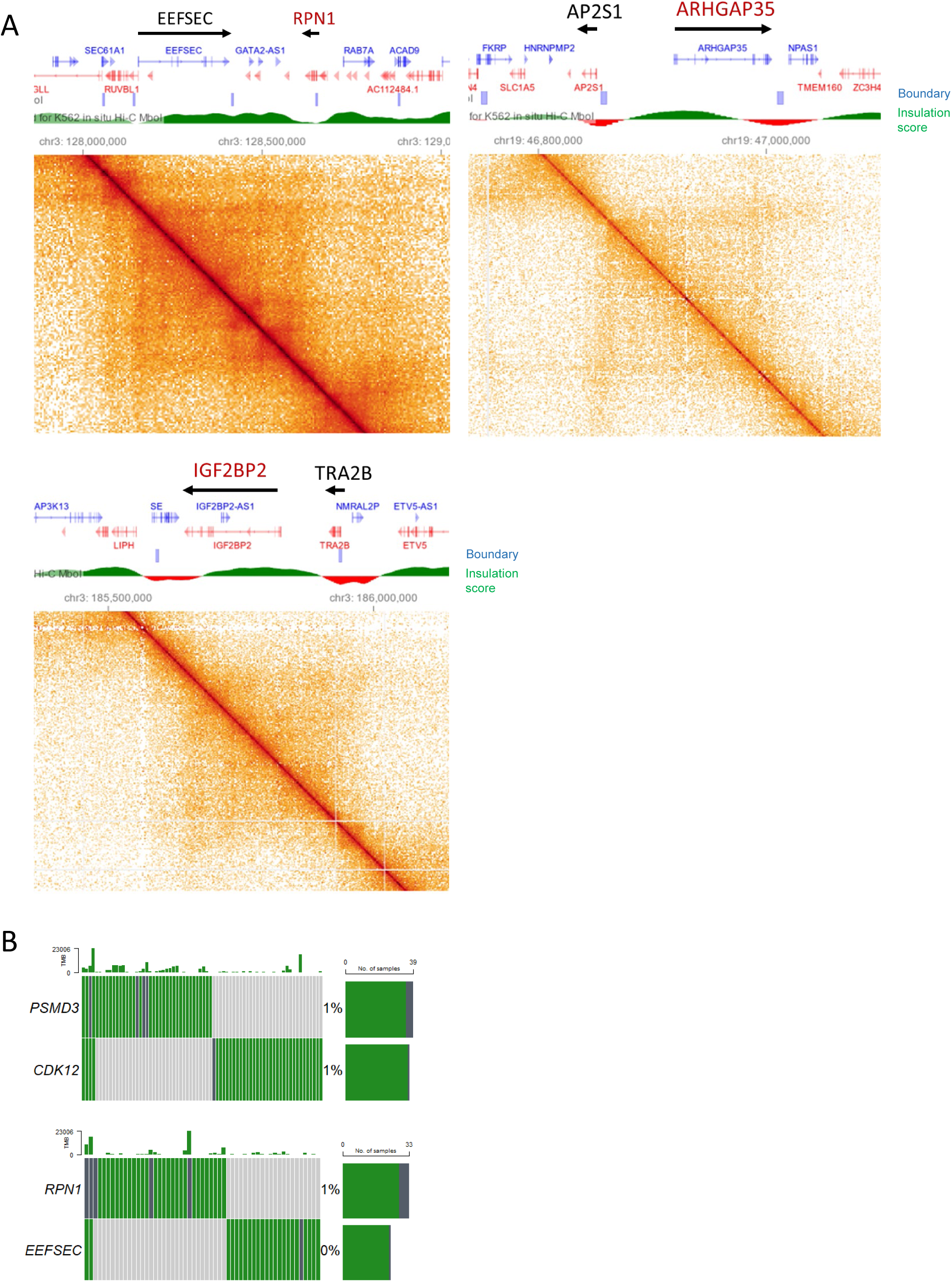
TAD boundary gene and TSS mutations. **A.**K562 HiC maps(8,34) of *EEFSEC*-*RPN1*, *AP2S1*-*ARHGAP35*, *IGF2BP2*-*TRA2B* pairs, insulation score and TAD boundary are shown above the HiC matrix. Names of oncogenes are in red. **B**. Oncoplots shown the landscape of ICGC mutations located in the promoters of TAD anchor non-oncogene-oncogene pairs, *PSMD3*- *CDK12* and *RPN1*-*EEFSEC*.

